# UPLC-ESI-MS based lipidomics revealed novel biomarkers in insulin receptor knockdown induced type 2 diabetes model of *Drosophila*

**DOI:** 10.64898/2026.08.20.745875

**Authors:** Prabhat Kumar, Zeeshan Fatima, Pradeep Kumar, Rohit Kumar, Brijesh Singh Chauhan, Saripella Srikrishna

## Abstract

Type 2 diabetes (T2D) is a prevalent metabolic disorder affecting millions worldwide, characterized by insulin resistance and impaired glucose homeostasis. While mammalian models are widely used, *Drosophila melanogaster* provides a powerful alternative due to its conserved insulin signaling pathways, genetic tractability, and suitability for high throughput studies. In addition to glucose dysregulation, lipid metabolism plays a crucial role in T2D pathophysiology, as alterations in lipid composition contribute to insulin resistance and metabolic dysfunction. Lipidomic studies have emerged as an essential approach to identify metabolic signatures and potential biomarkers for disease progression and therapeutic targeting. In this study, T2D like model was established by inducing insulin resistance through knockdown of the insulin receptor in brain insulin-producing cells using the *dilp2-Gal4>UAS-InR^RNAi^*system. This genetic manipulation resulted in significant metabolic dysregulation, including elevated glucose, trehalose, and triacylglyceride levels, along with increased oxidative stress indicators. Additionally, mRNA expression analysis of key insulin signaling components, including insulin receptor substrate 1, *dilp2*, *dilp3*, *dilp5*, and phosphorylated Akt, further validated the model. To further investigate metabolic alterations, Lipid profiling was performed using ultra-performance liquid chromatography coupled with quadrupole time-of-flight mass spectrometry (UPLC-QTOF-MS) in non targeted LC-MS-based metabolomics approach to identify lipid biomarkers associated with T2D. Multivariate statistical analyses, including PCA and PLS-DA, revealed distinct lipid signatures between wild-type and T2D flies. Notably, specific phosphatidylglycerol species PG 34:0, PG 34:4, PA 38:3, PIP 38:1, PIP2 38:6, and LPS 24:0 demonstrated an area under the curve (AUC) of 1, indicating their strong reliability as lipid biomarkers for T2D diagnosis.

**Research Highlights:**

- *InR* was knocked down in *Drosophila* IPCs to T2D like model.
- T2D flies showed hyperglycemia, elevated lipids, and altered dilp2, 3, and 5 levels
- Insulin signaling was impaired, with increased oxidative stress and pAkt levels
- Six novel lipid biomarkers of T2D were identified via UPLC-ESI-MS lipidomics
- PG 34:0, PG 34:4, PA 38:3, PIP 38:1, PIP2 38:6, LPS 24:0 are potential T2D biomarkers in flies

## 1. Introduction

Type 2 diabetes (T2D) represents a significant global health issue and is the leading cause of mortality associated with chronic diseases worldwide. In 2019, an estimated 463 million individuals globally were diagnosed with T2D, with forecasts anticipating that this figure may escalate to 700 million by 2045[1]. The two primary approaches are used to establish T2D models in *Drosophila,* first dietary induction through a high sugar diet, and second genetic manipulation via knockdown of conserved genes involved in the insulin signaling pathway include.[2, 3]. The Insulin producing cells (IPCs) in the *Drosophila* central nervous system function analogously to pancreatic β-cells in humans. The insulin signaling pathway is highly conserved across *Drosophila* and mammals, playing comparable roles in metabolic regulation [4–6]. The median neurosecretory cluster (NSC) of the *Drosophila* brain harbors IPCs. These cells are responsible for the secretion of three out of the eight *Drosophila* insulin like peptides (DILPs), specifically *dilp2*, *dilp3*, and *dilp5*, which play a critical role in the regulation of metabolic processes [7, 8]. In *Drosophila*, insulin/insulin-like growth factor signaling is mediated through the insulin receptor (InR), a conserved receptor tyrosine kinase. Upon ligand binding, *InR* activates downstream adaptor proteins, including Chico, the principal IRS homolog in Drosophila, as well as Lnk, a member of the SH2B family. These adaptor proteins facilitate activation of the phosphoinositide 3-kinase (PI3K)/protein kinase B (Akt) pathway. PI3K generates phosphatidylinositol-3,4,5-trisphosphate (PIP3), which promotes the recruitment and activation of Akt and phosphoinositide-dependent kinase-1 (PDK1). This signaling cascade regulates key physiological processes, including growth, metabolism, energy homeostasis, and longevity [9, 10]. Together with phosphoinositide-dependent kinase-1 (PDK1), Akt is activated through phosphorylation, generating phosphorylated Akt (p-Akt)[11, 12]. Activated p-Akt subsequently phosphorylates downstream targets, including glycogen synthase kinase-3β (GSK-3β), thereby inhibiting its activity and promoting the downstream effects of insulin signaling [13, 14]. The T2D is reportedly significantly correlated with oxidative stress, and sustained chronic elevated glucose can result in excessive ROS production in the context of T2D [15, 16]. The organism possesses a highly conserved antioxidant defense system that includes enzymatic components such as superoxide dismutase (SOD) and catalase (CAT), as well as non-enzymatic pathways. SOD is essential for the dismutation of superoxide radicals, thereby limiting oxidative stress. In T2D, hyperglycemia disrupts this defense mechanism, leading to reduced SOD activity. This decline underscores the enzyme’s pivotal role in counteracting glucose induced oxidative damage and preserving cellular homeostasis [17]. The vital antioxidant enzyme catalase protects β-cells from oxidative stress induced damage linked to ROS in T2D by catalysing the conversion of hydrogen peroxide into oxygen and water [18].

Metabolomics involves the application of comprehensive methods to identify small molecules, metabolites, and metabolic profiles that reveal the cellular response phenotype to genetic alteration or pathophysiological triggers[19]. Metabolomics provides valuable opportunities to develop biomarkers for diagnosing diseases, predicting their progression, and optimizing treatments. The identification of novel biomarkers and pathways holds the potential to enhance our comprehension of the pathophysiological alterations associated with T2D [20, 21]. Lipidomics, a specialized domain within metabolomics, concentrates on the systematic examination of the entire lipid repertoire, referred to as the lipidome, present in any biological system. We implemented UPLC-MS to perform lipid metabolomics on entire *Drosophila* specimens, with the objective of investigating metabolic patterns and prospective biomarkers within a T2D model, thus augmenting our understanding of the pathological transformations associated with the condition [22, 23]. Our goal was to ascertain lipid molecular differences between the two cohorts and to evaluate selected biomarkers for the prediction of T2D.

## 2. Material and Methods

### 2.1. *Drosophila* culture and husbandry

Flies stocks, including *dilp2-GAL4-GFP/CyO*, *UAS-InR ^RNAi^*, and *UAS-GFP*, were obtained from the Bloomington *Drosophila* Stock Center (BDSC), Indiana, USA, and the National Institute of Genetics (NIG), Japan. To generate the insulin receptor (InR) knockdown T2D like model, virgin females carrying *UAS-InR ^RNAi^* were crossed with males carrying the *dilp2-GAL4-GFP/CyO* driver. The F1 progeny expressing both *dilp2-GAL4-GFP > UAS-InR^RNAi^* were selected and used as the T2D-like model.

For the wild-type (WT) control, *dilp2-GAL4-GFP/CyO* flies were used. All fly stocks and experimental groups were reared on standard cornmeal agar medium and maintained in a temperature-controlled BOD incubator at 24 ± 1°C under a 12 h light/12 h dark photoperiod.

### 2.2 Measurement of adult body length and area of flies

Morphological images of male and female *Drosophila* were acquired using a Dewinter Trinocular zoom microscope. The body length and body area defined from the anterior margin of the head to the posterior end of the abdomen were quantified for each sex using ImageJ software for image analysis.

### 2.3 Circulatory glucose, trehalose, and triglyceride quantification

Third instar larvae (n = 15) were collected, rinsed with 1X PBS, and subjected to hemolymph extraction. The extracted hemolymph was then analyzed using previously described methodologies. Glucose levels were quantified using a glucose (hexokinase) assay kit (Cat# GAHK20, Sigma-Aldrich). To quantify trehalose, hemolymph was enzymatically treated with porcine trehalase (Cat# T8778) Sigma, absorbance was measured at 340 nm [24, 25]. The larvae were homogenized in 150 µL lysis buffer (0.05% PBST), incubated at 75°C for 10 min, and centrifuged (12,000 rpm, 10 min, 4°C). A 20 µL supernatant aliquot was mixed with triglyceride assay reagent, incubated at 37°C for 15 min, and absorbance was measured at 505 nm using Biotek plate reader.

### 2.4 Oxidative stress analysis

*Drosophila* strains (n = 10) were sourced from each respective group, including WT and T2D, for the assessment of H2O2 levels, SOD activity, catalase enzyme activity, and LPO levels. Furthermore, the quantification of H2O2 was conducted utilizing the methodology established [26]. The activity of SOD was quantified by measuring the 50% inhibition of the chromogenic compound nitro blue tetrazolium (NBT) after 30 minutes of exposure to light, [27]. Catalase activity was assessed following the protocol outlined [28]. The lipid peroxidation (LPO) was evaluated under standardized conditions utilizing thiobarbituric acid reactive substances (TBARS), as per the methodology described[29].

### 2.5 RNA extraction, cDNA preparation, and qRT-PCR quantification

flies heads (n = 90) was decapitated throught the fine needles of both groups. The biological sample was homogenized with 500 µl of TRIzol reagent (Sigma Lot# BCCF 8996), A protocol recommended by the manufacturer was followed to isolate the total RNA. A nanodrop spectrophotometer determined the RNA concentration, and cDNA samples were synthesized using (1000ng) extracted RNA through reverse transcriptase[30]. cDNA was used for qPCR to quantify gene expression, normalized to rp49, and analyzed by the ΔΔCt method with gene-specific primers.

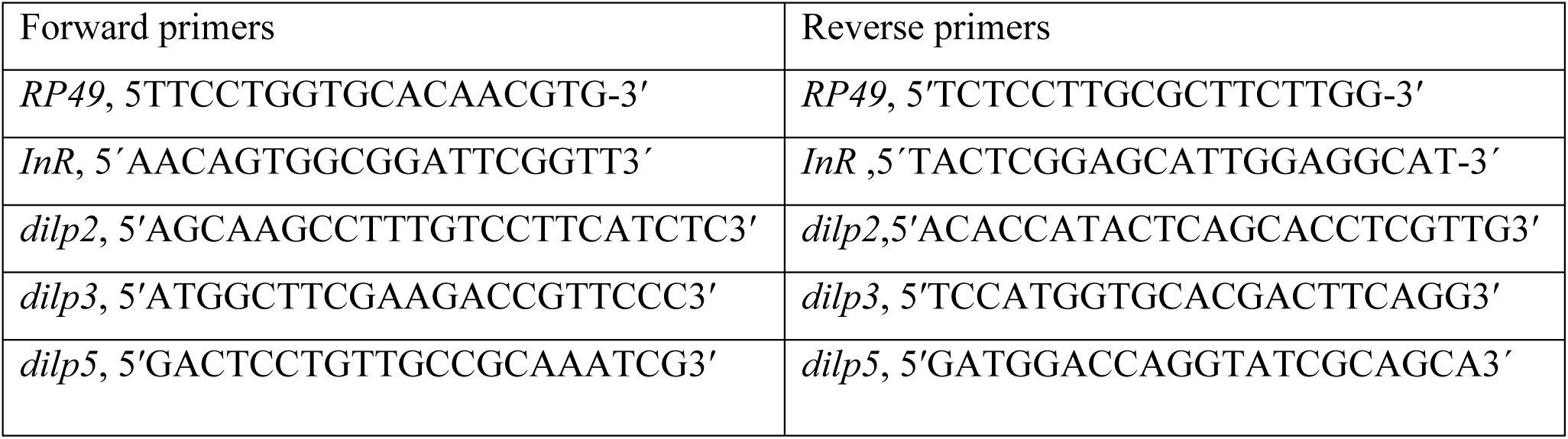

### 2.6 Protein isolation and western blot analysis

Protein was isolated from fly head of WT and T2D, in three biological replicates, and SDS-PAGE, western blotting, and immunodetection were completed as per the protocol described by Kumar et al [31]. The pAkt (Thr 342) polyclonal antibody (Cat# PA5 95699 1:7500) and E7 anti-beta tubulin (DSHB 1:7500). The secondary antibodies utilized were sourced from DSHB, including Goat anti-rabbit IgG (Cat# 0600580051730, 1:10000) and Goat anti-mouse IgG (Cat# 0600680251730, 1:10000). The blots were incubated with Goat anti-Rabbit or Goat anti-mouse IgG antibodies conjugated to horseradish peroxidase and diluted to a concentration of 1:10000. Detection of the blots was performed utilizing the enhanced chemiluminescence methodology of ClarityTM Western ECL substrate (BIO-RAD Cat# 170-5060), with the blot images captured using the ChemiDocTM XRS+ imaging system (BIO-RAD)Furthermore, blot intensity analysis with β-tubulin normalization was performed using Quantity One software.

### 2.7 Lipid extraction and UPLC-Q-TOF-MS analysis

Lipids were extracted from male and female flies of both groups in biological replicates. Further, lipid profiles were characterized using UPLC-Q-TOF-MS as per the method suggested by Islam et al [32]. WT and T2D genotypes were used for the lipid extraction study. Three biological replicates were taken for each group for lipid extraction. Lipid extraction was done according to the previously described Folch method [39]. Briefly, the flies were washed with 3× ice-cold PBS (pH 7.4). The adult flies were crushed and homogenized in an aqueous solution for 3 min and suspended in chloroform and methanol in the ratio of (1:2). Suspensions were shaken well and centrifuged at 5000× g at 4 ◦C for 15 min. The supernatant was transferred to another glass vial and then the remaining chloroform was added and filtered through Whatman No. 1 filter paper. The extract was then washed with 0.88% KCl to remove the non-lipid contamination. The lower dense layer of chloroform containing lipid was taken by glass Pasteur pipette in a 5 mL glass vial with Teflon capping. The extraction solvents were evaporated using N2 gas flux and stored at −20 ◦C until further use.

### 2.8 Ultraperformance Liquid Chromatography-Electrospray Ionization-Mass Spectrometry (UPLC-ESI-MS)

To perform reverse-phase ultrahigh pressure liquid chromatography (UHPLC, Exion LC Sciex, Framingham, MA, USA) using ZORBAX RRHD Eclipse Plus C18 column with 1.8 μ particle size, 2.1 mm inner diameter, and length 100 mm were purchased from Agilent Technologies, Santa Clara, CA, USA coupled to a hybrid triple quadrupole/linear ion trap mass spectrometer (4500 Q-TRAP, SCIEX, Foster, CA, USA). The dried sample was reconstituted in mobile phase solvent B before use. The sample was introduced using an autosampler with 5 mL of sample injection volume at a 0.1 mL/min flow rate. The elution was done for 30 min, using mobile phases A and B, solvent A has 10% methanol and 90% buffer of ammonium acetate (5 mM) dissolved in water and acetonitrile (95:5), and solvent B has 5% isopropanol, 10% methanol, and 85% acetonitrile. All the solvents used were MS grade and purchased from Honeywell. The source temperature was 120◦C, the desolation temperature was 350◦C, and the cone voltage was set at 40 volts (V). Capillary voltage (i.e., spray voltage) was set to 3.50 kilovolts (kV). The data were recorded in the mass range m/z 200–2000 Da in-electrospray ionization (ESI) positive and negative ion mode. The data processing was done with the Analyst software, where each chromatogram was smoothened and background was subtracted. Technical triplicate LC-ESI-MS datasets were acquired and extent of reproducibility of the data was verified. Further analysis was carried out with the reproducible LC-ESI-MS data and only those peak m/z values in the mass spectra have been considered for interpretation, whose intensities were higher than 5000 (namely, the threshold value to ignore the peaks with poor signal-to-noise ratios). Analysis of the acquired data was performed by two different software viz. Lipid View and mass spectrometry-based lipid (ome) Analyzer and molecular platform (MS-LAMP) software. MS-LAMP software is a graphical user interface (GUI) standalone program built using Perl: Tk, based on lipid metabolites and pathways strategy consortium (LIPID MAPS; www.lipidmaps.org). Further, it needs to be noted that the data herein were analyzed qualitatively only. All the experiments were performed in biological duplicates with technical triplicates to ensure reproducibility and accuracy.

Data were processed and uploaded into MetaboAnalyst 5.0 (http://www.metaboanalyst.ca) and normalized for uni and multivariate statistical analysis in both positive as well as negative modes separately. The initial exploratory data analysis was done by univariate one-way analysis of variance (ANOVA). The unsupervised PCA method was applied through the prcomp package and the calculation was based on singular value decomposition to explain the variance in the data set. Then, the partial least square (PLS) supervised method using multivariate regression technique was applied using the plsr function provided by rpls package. The classification and cross-validation were done using their corresponding wrapper function through caret package. The sparse PLS-DA (sPLD-DA) algorithm was used to effectively reduce the number of variables.

The ROC curve was created through the MetaboAnalyst R by computing ratios between all possible metabolite pairs. Three algorithms (PLS-DA, SVM, RF) were used to evaluate and validate the lipid species for biomarker search. Lipids species for biomarker search were selected based on kinds of literature and overall ranks obtained by AUC, t-statistic, or fold-change.

### 2.17 Statistical Analysis

All the data were conducted in biological triplicates. The data were expressed as mean ± standard error mean (SEM). Statistical analysis was performed using t-test in Graph Pad Prism 8 Data were considered significant between groups at p < 0.05.

## 3.0 Results

### 3.1 Knockdown of insulin receptors in IPC cells in the brain of flies changed the phenotype

To evaluate the effect of insulin receptor (*InR*) knockdown in insulin-producing cells (IPCs), we performed immunofluorescence imaging of third instar larvae brain lobe of flies using GFP as a reporter. In WT flies, strong GFP expression was detected in the median neurosecretory cells of the brain, representing intact IPC activity (Figure a). In contrast, T2D model flies exhibited markedly reduced GFP fluorescence (Figure a2), indicating significant suppression of insulin signaling. Bar line intensity profiles (Figure 1 a1 and a3) further confirmed a substantial decline in GFP intensity in T2D larvae brain relative to WT. In females (Figure 1b), T2D flies displayed visibly smaller body size compared to WT. Quantitative measurements revealed a significant reduction in body weight (Figure 1 b1), body length (Figure 1 b2), and total body area (Figure 1 b3) in T2D females. Similarly, male T2D flies (Figure 1 c) exhibited smaller than WT counterparts, with significant decreases in body weight (Figure 1c1), length (Figure 1 c2), and area (Figure 1 c3).

**Figure 1.**
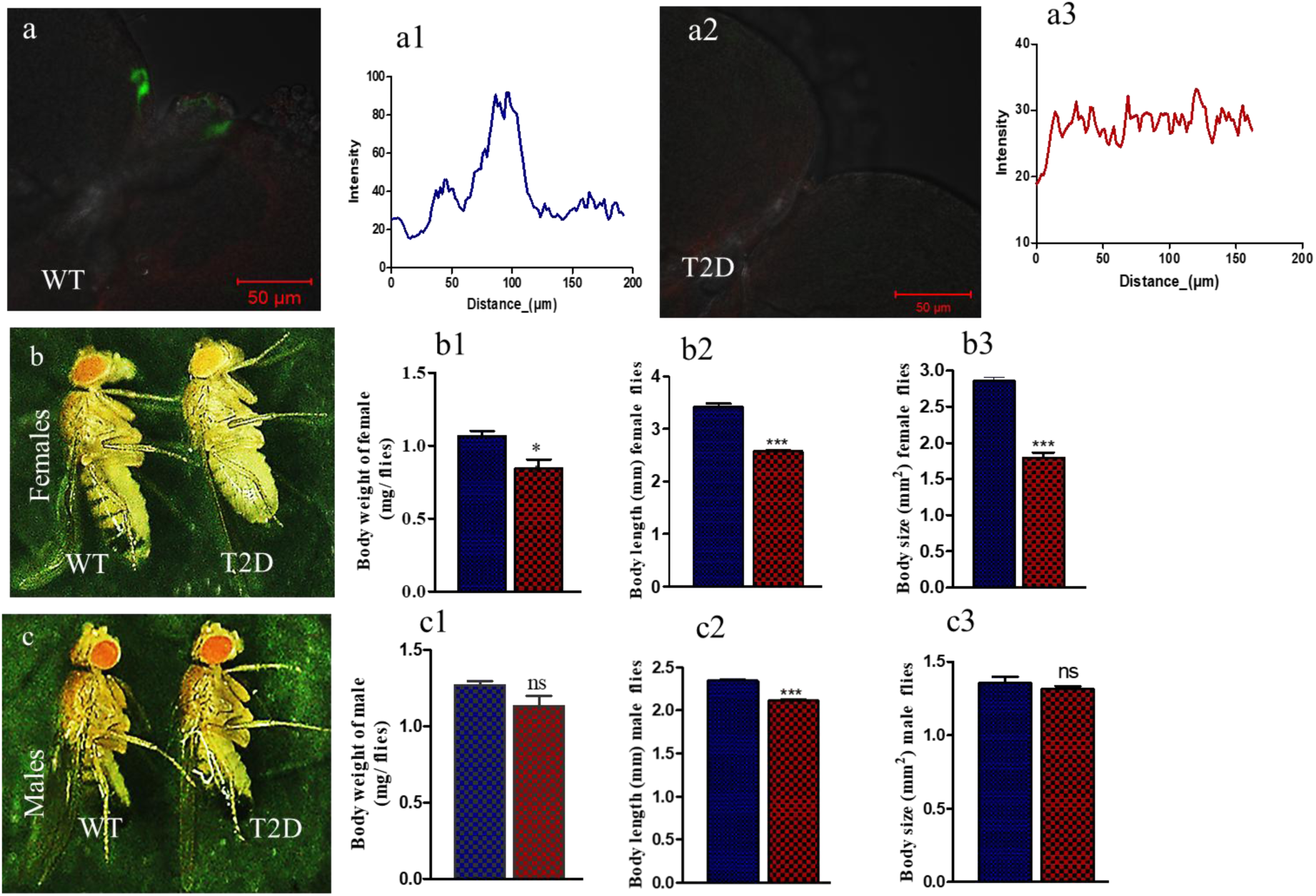
Illustrates the phenotypic alterations observed in a T2D compared to WT. (a) WT third instar larvae of brain lobe and (a1) intensity, (a2) T2D third instar larvae brain lobe and (a3) intensity, female represent (b),weight (b1), length (b2),and area (b3). male flies represent (c), weight (c1), length (c2), and area (c3) (n = 10) in triplicate of significantly decreased a. **p<0.0082, b. nsp>0.0701 c. ***p<0.0003, d.ns p>0.3902 e. ***p<0.0002, f. ***p<0.0006.

### 3.2 Alterations in Glucose, Trehalose, and Triglyceride Levels in T2D third instar larvae

We assessed glucose concentrations in the circulating hemolymph of WT and T2D third instar larvae. The glucose levels in WT larvae were measured at 6 mg/dl per third instar larvae, while T2D larvae exhibited a significantly higher concentration of 14.13 mg/dl per larvae. This corresponds to a 2.34-fold increase in glucose levels in T2D third instar larvae compared to the WT (Figure 2 a). We have also measured the trehalose levels in WT and T2D samples. Notably, we observed a 1.52-fold elevation in trehalose levels in T2D compared to WT (Figure 2 b).

**Figure 2.**
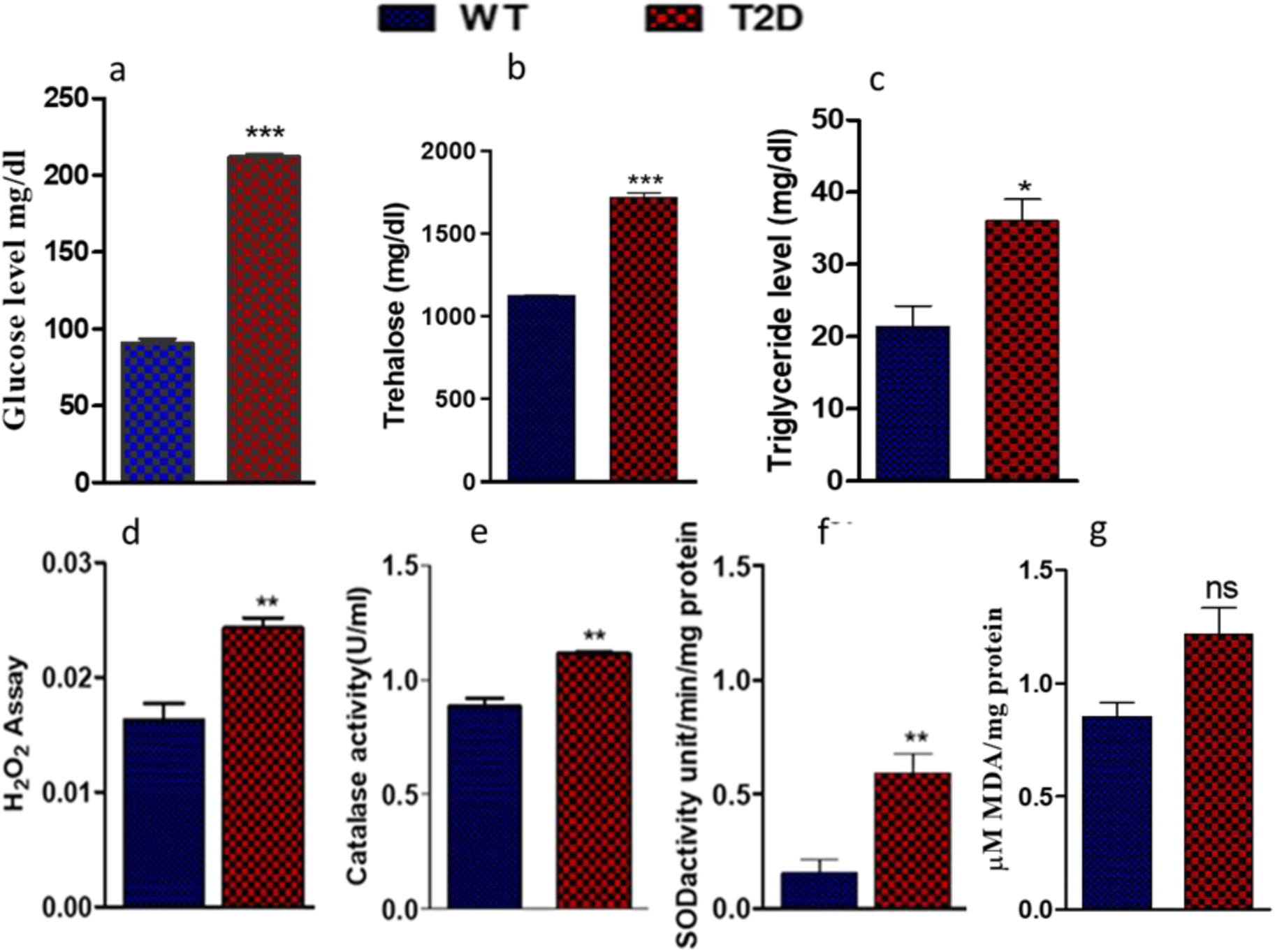
Represent oxidative enzyme in the case of control and T2D flies, a. glucose b. trehalose, c. triacylglyceride, from the hemolymph of (n = 15) third instar larvae, d. H2O2 assay, e. catalase activity, f. SOD assay and g. MDA from the flies. statistical significance assessed and p-values reported accordingly **p< 0.007, **p< 0.003, *p< 0.033, nsp >0.052, and ***p< 0.001.

Additionally, we assessed triglyceride levels in third instar larvae from both WT and T2D groups. Remarkably, triglyceride levels were significantly increased by 1.71-fold in T2D third instar larvae compared to WT (Figure 2 c)

### 3.3 Assessment of Oxidative Stress

Our analysis revealed a significant elevation in oxidative stress markers in T2D *Drosophila* compared to WT controls. The H₂O₂ levels were 1.5-fold higher in T2D flies than in WT (Figure 2 d). Further, SOD activity was markedly elevated in T2D flies, showing a 3.83-fold increase compared to WT (Figure 2 e). Similarly, catalase activity, measured as U/ml of protein, increased from 0.882 U/ml in WT to 1.114 U/ml in T2D, representing a 1.263-fold increase (Figure 2 f). The MDA level was increased to 1.31-fold T2D as compred to WT (Figure 2 g).

### 3.4 Quantitative Assessment of Gene Expression

We have found the relative mRNA expression of the *InR* gene in T2D flies was significantly reduced 2.5-fold compared to WT (Figure 3 a). In T2D flies, *dilp2, dilp3*, and *dilp5* were upregulated compared to WT. Specifically, in T2D dilp2 expression increased 1.5 fold (Figure 3 b), *dilp3* exhibited a 5.01-fold increase (Figure 3 c), and *dilp5* expression was 2 -fold higher relative to WT flies (Figure 3 d).

**Figure 3.**
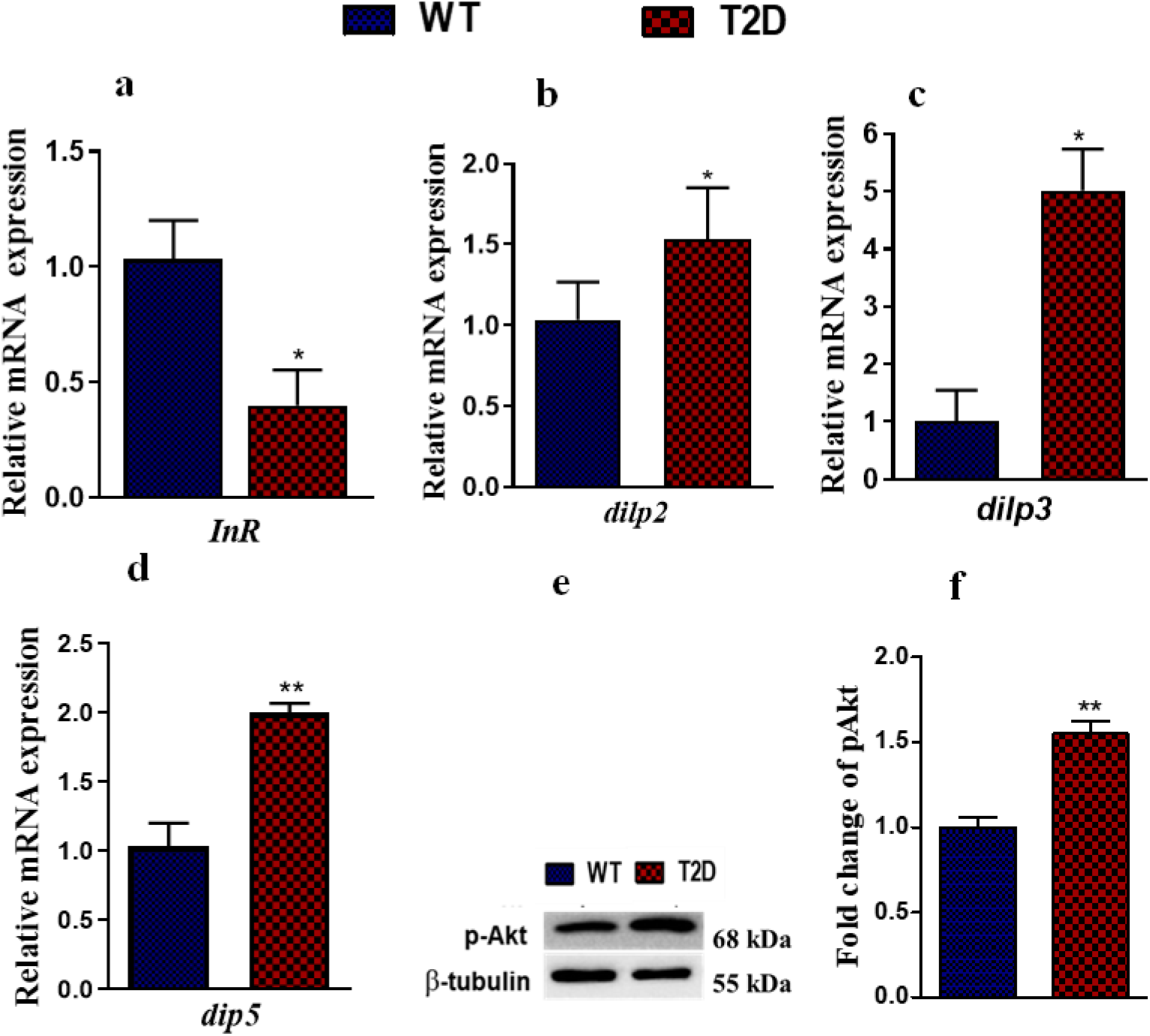
The insulin receptor was knocked down in insulin-producing cells within the adult brain (n = 80). We assessed the expression levels of each gene using quantitative real-time PCR (qRT-PCR) of a. *InR*, b. *dilp2*, c. *dilp3*, and d. *dilp5*, where p values were considered significant at p<0.05. Additionally, e. pAkt blot and f. pAkt fold change in WT and T2D, analyzed using Quality One (BioRad), with significance at **p < 0.004.

### 3.5 The levels of pAkt protein expression

Western blot analysis revealed a marked increase in phosphorylated pAkt levels in type T2D compared to WT flies. Densitometric analysis revealed an approximately 1.5-fold increase in p-Akt expression in the T2D group compared with the WT group (Figure 3 f).

### 3.6 Multivariate lipid profile

To evaluate the global lipidomic alterations associated with T2D in *Drosophila*, we conducted Principal Component Analysis (PCA) on the lipid datasets acquired in both +ESI) and negative (–ESI) electrospray ionization modes. The scree plots (Figure 4 A a and e) illustrate the distribution of total variance across principal components, where the green line represents cumulative variance and the blue line depicts individual signal variance for each principal component (PC). In +ESI mode, PC1 and PC2 accounted for 46.3% and 20.8% of the total variance, respectively, while in –ESI mode, PC1 and PC2 explained 48.4% and 24.2%, respectively. These high variance values suggest that first two PCs capture the majority of differences in lipid profiles between the experimental groups. Two dimensional (2D) and three-dimensional (3D) PCA score plots (Figure 4A b, f and c, g) revealed clear separation between WT and T2D flies in both ionization modes, with minimal overlap in sample clustering. This indicates substantial shifts in lipid composition and abundance associated with the T2D state.

**Figure 4.**
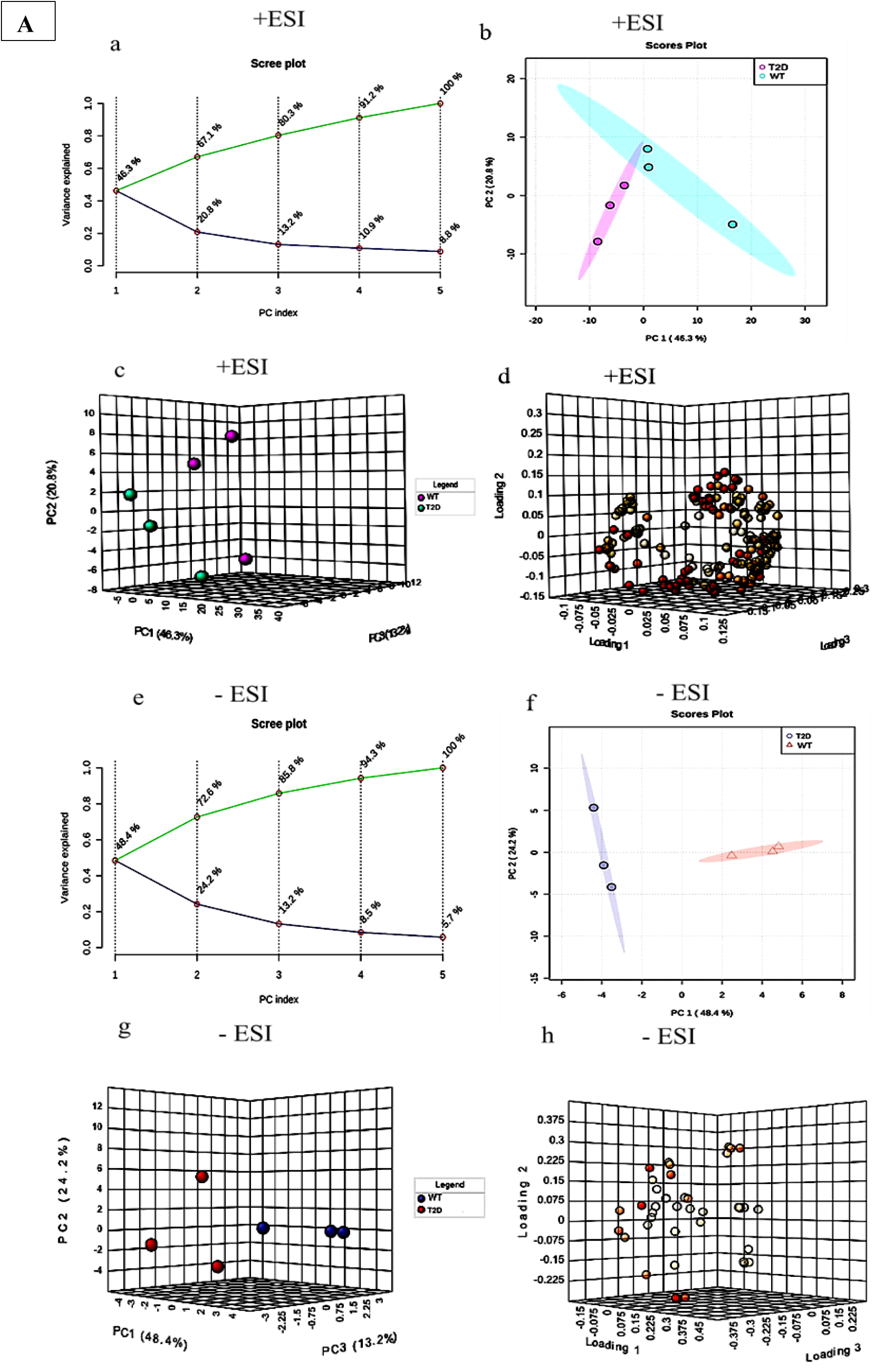
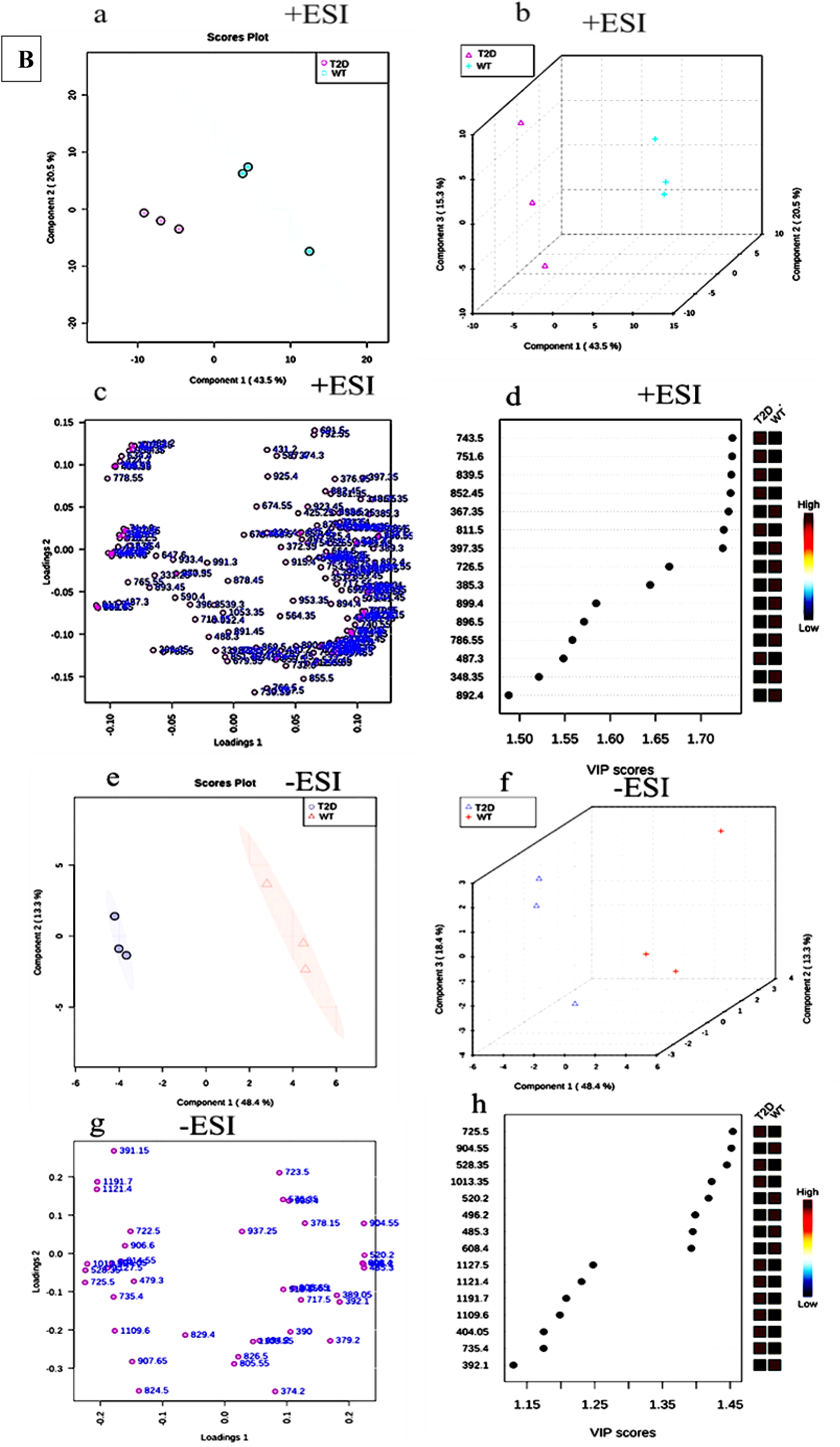
(A) The experimental data were analyzed using PCA, with the results illustrated in a scree plot showing the total variance (green line). The variance explained by each principal component is depicted by signal variance (blue line) for each PC index (a & e). Additionally, the two and three dimensional score plots (b & f and c & g), as well as the loading plots (d & h), were generated for both +ESI and -ESI. (B) PLS-DA of two- and three-dimensional score plots (a & e and b & f), loading plots (c & g), and VIP scores (d & h) were created for +ESI and -ESI.

Furthermore, the corresponding PCA loading plots (Figure 4(A) d and h) highlight the lipid species that most strongly contribute to the observed variance and group separation. To further refine group separation and identify the most influential lipid species contributing to the T2D phenotype, Partial Least Squares Discriminant Analysis (PLS-DA) was performed under both +ESI and –ESI ion modes. The two and three dimensional PLS-DA score plots (Figure 4B a, b for +ESI and e, f for –ESI) demonstrated clear discrimination between T2D and WT groups. The tight clustering and minimal overlap observed in these supervised models confirm a high degree of metabolic distinction between the groups.

The PLS-DA loading plots (Figure 4B c for +ESI and g for –ESI) visualize the individual lipid variables that contribute most significantly to the group separation. The spread and clustering of lipids in the loading space reflect the metabolic complexity and variation specific to T2D induced lipid alterations. Notably, several lipid ions cluster in defined regions, suggesting that select lipid subclasses (e.g., phosphatidylglycerols, phosphoinositides, and lysophospholipids) play central roles in the observed metabolic shifts. Variable Importance in Projection (VIP) scores were calculated to quantitatively rank the discriminatory power of individual lipid features. The VIP plots (Figure 4B d for +ESI and h for –ESI) highlight the top ranked lipids that most strongly influence the classification between T2D and WT samples. The heatmap bars adjacent to VIP markers visually indicate the relative abundance of these ions across groups, providing additional biological context to the statistical relevance.

Together, these PLS-DA results reinforce the conclusions drawn from PCA showing that T2D in *Drosophila* is associated with a robust and distinct lipidomic shift. Moreover, the identification of specific VIP scored lipid markers provides a focused list of candidate biomolecules for further functional validation and mechanistic investigation. Using MS LAMP, we classified and characterized lipid distributions, identifying over 2,500 lipid molecules across all lipid classes in both +ESI and -ESI modes (Table S1). The Variable Importance in Projection (VIP) scores, derived from the Partial Least Squares Discriminant Analysis (PLS-DA), were used to identify the top 15 fatty acids based on peak intensity, providing a prioritized list of lipid molecules with the greatest discriminatory power between WT and T2D flies (Figures 4B d & h).

### 3.7 Univariant lipid profile

We performed a comprehensive lipid analysis of WT and T2D *Drosophila* strains using univariate and multivariate approaches to assess alterations in lipid composition between the two groups. To quantify the differences, fold change (FC) was calculated for the intragroup concentrations of free fatty acids, with positive values indicating higher fatty acid levels in T2D and negative values indicating lower concentrations in WT. The fold change analysis was conducted in both both +ESI and -ESI ion.

In +ESI, a total of 44 lipids were analyzed. Among these, 20 lipids exhibited increased levels, while 24 showed decreased levels in T2D compared to WT (Table S7). Similarly, in -ESI ion mode, 18 lipids were identified, with 11 showing increased levels and 7 demonstrating decreased levels in T2D flies (Table S7). Statistical analysis was carried out using a t-test based on fold change, revealing significant alterations in lipid profiles. In positive ion mode, 34 lipids were significantly altered, with 23 downregulated and 11 upregulated. In -ESI ion mode, 16 lipids showed significant changes, with 9 downregulated and 7 upregulated (Tables S8, S9). To investigate lipidomic alterations associated with T2D in *Drosophila*, we performed untargeted UPLC-ESI-MS analysis in both positive (+ESI) and negative (−ESI) ion modes, followed by volcano plot based differential analysis and hierarchical clustering. In the +ESI mode, volcano plot analysis (Figure 5 a) identified 26 significantly dysregulated metabolites (Table S10), with several ions showing strong upregulation in T2D compared to WT. Notably, ions at m/z 743.5, 751.6, 839.5, 852.45, and 811.5 exhibited high fold changes (log₂FC > 2.2) and strong statistical significance (p < 0.0001). In contrast, ions such as m/z 367.35, 397.35, and 726.5 were significantly downregulated (log₂FC < −2.5). These features represent prominent lipid changes in the diabetic condition. Similarly, in the −ESI mode (Figure 5 b), 15 key ions were identified (Table S11), including strongly upregulated features such as m/z 725.5 (log₂FC = 3.29), 528.35 (log₂FC = 3.07), and 1013.35 (log₂FC = 2.97), along with downregulated ions including m/z 904.55, 520.2, and 485.3, all showing log₂FC values below −1.8. These alterations reflect significant metabolic reprogramming associated with impaired insulin signaling.

**Figure 5.**
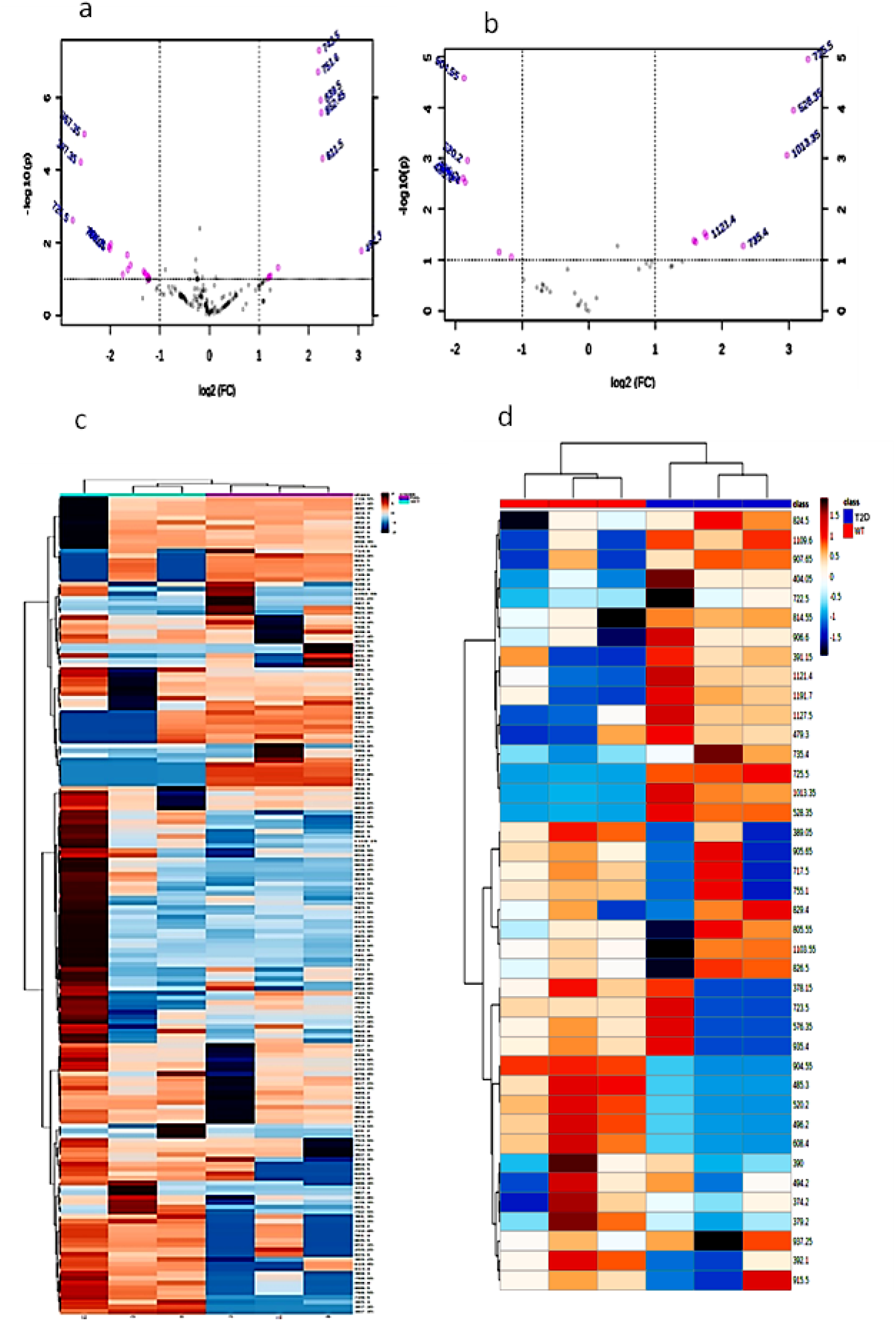
The volcano plot displays the differentially expressed metabolites between WT and T2D flies of both +ESI and -ESI ions modes (a & b). Heat map visualizations show the fatty acid content, with WT indicated in red and T2D in blue (c & d).

To further explore these differences, hierarchical clustering heatmaps were generated based on the top discriminatory features in both ionization modes. In +ESI mode (Figure 5 c), samples clustered distinctly by genotype, with widespread changes in lipid abundance between WT and T2D flies. Similarly, the −ESI mode heatmap (Figure 5 d) demonstrated clear separation of T2D and WT samples, with several metabolites contributing prominently to group discrimination.

### 3.8 Differential free fatty acids related to metabolic disorder

To identify potential lipid biomarkers distinguishing T2D from WT *Drosophila*, targeted lipidomic profiling followed by receiver operating characteristic (ROC) analysis was performed. A subset of seven lipid species displayed an area under the ROC curve (AUC) of 1.0, indicating perfect discriminatory power between T2D and WT groups (Figure 6 a–g). These lipids were further validated by comparing normalized expression profiles and raw intensity values (Tables S4 and S5). In the positive electrospray ionization (+ESI) mode, phosphatidylglycerol species PG 34:0 (m/z 751.6) and PG 34:4 (m/z 743.3) were completely absent in all WT biological replicates but were highly expressed in T2D flies, with raw intensities averaging approximately 1.7 million (Table S4). Similarly, phosphatidic acid (PA) 38:3 (m/z 725.5) was undetectable in WT but markedly elevated in T2D samples (average intensity ≈ 131,828), indicating significant upregulation of these glycerophospholipids under diabetic conditions (Table S5). In contrast, phosphoinositide species including phosphatidylinositol phosphate (PIP) 38:1 (m/z 485.3) and phosphatidylinositol bisphosphate (PIP₂) 38:6 (m/z 520.2) showed the opposite pattern, with high expression in WT (averaging ∼181,698 and ∼291,838, respectively) but complete absence in T2D flies. Additionally, lysophosphatidylserine (LPS) 24:0 (m/z 608.4) was robustly detected in WT (average ∼281,087) and completely depleted in diabetic flies. The absence or exclusive presence of these lipid species between the two groups resulted in zero overlap and a perfect classification, reflected by an AUC value of 1.0 across all ROC analyses. These lipid species serve as robust and non-overlapping biomarkers for metabolic dysfunction in this *Drosophila* T2D model. In +ESI mode, PG 34:0 and PG 34:4 (Figure 6 a and b) showed significantly increased levels in T2D flies compared to WT. In −ESI mode, PA 38:3 (Figure 6 c) was notably upregulated in T2D flies, indicating altered mitochondrial dynamics, including potential activation of fission and mitophagy processes. In contrast, PIP 38:1 and PIP2 38:6 (Figure 6 d and e) were significantly downregulated, pointing toward disruptions in phosphoinositide mediated insulin signaling, calcium homeostasis, and ER mitochondrial contact site function. Additionally, LPS 24:0 (Figure 6 f) levels were markedly decreased in the T2D group, suggesting increased lysophosphatidylserine accumulation, possibly reflecting lipotoxic stress and compromised mitochondrial membrane integrity.

**Figure 6.**
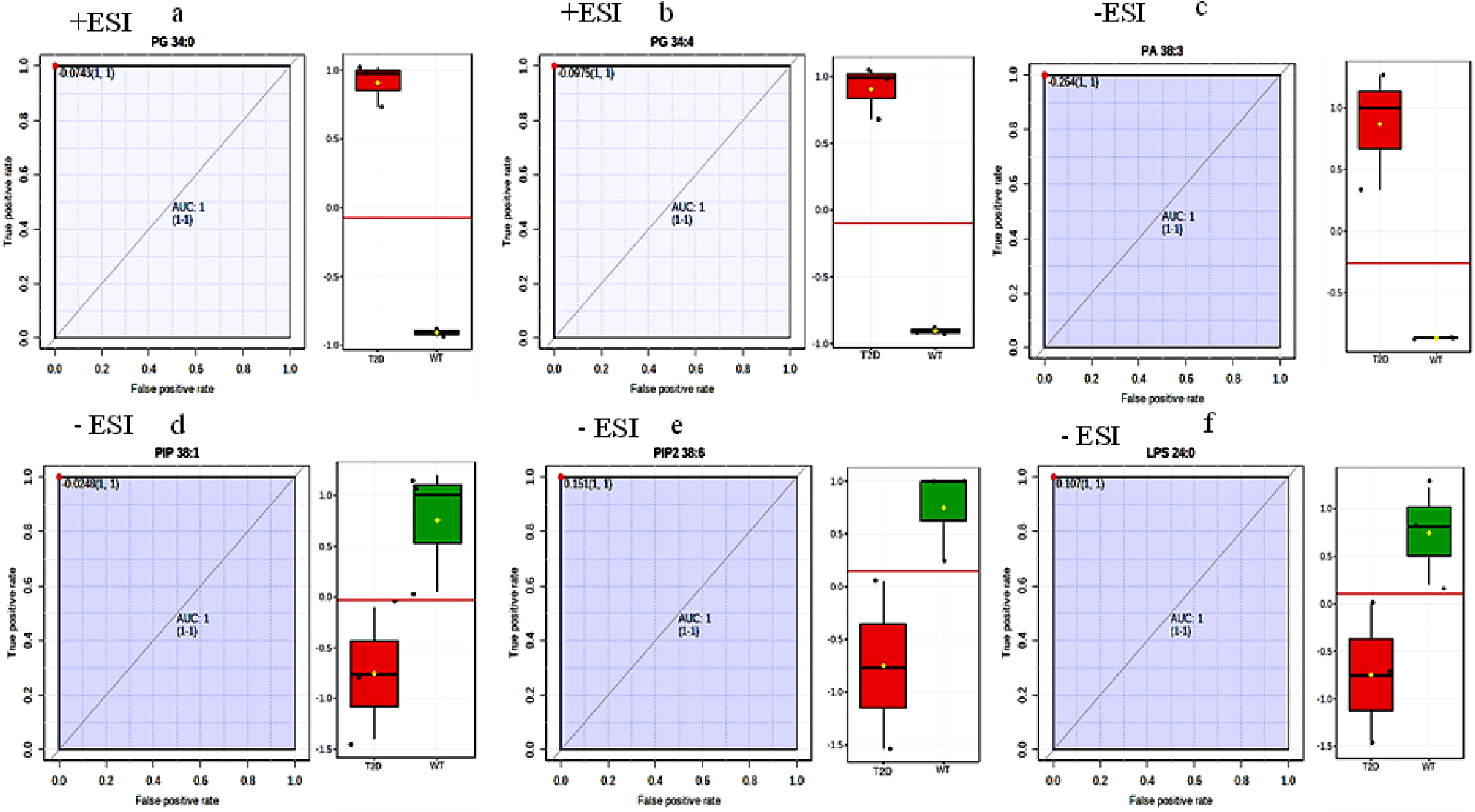
The six most discriminative lipid molecules identified through analysis of AUC, t-test, and fold change in both +ESI and -ESI ion modes for WT and T2D were: (a.) PG 34:0, (b.) PG 34:4, (c.) PA 38:3, (d.) PIP 38:1, (e.) PIP2 38:6, and (f.) LPS 24:0.

### 3.9 Lipidomic analysis reveals major lipid classes variation

We performed a comparative lipidomic profiling of WT and T2D *Drosophila* samples using mass spectrometry under both +ESI and -ESI ionization modes (Table S1). In the +ISE (Figure 7 a), the predominant lipid classes detected were glycerophospholipids and glycerolipids, followed by polyketides and sterol lipids. Notably, WT samples exhibited significantly higher abundance of glycerophospholipids and glycerolipids compared to the T2D samples, suggesting perturbations in membrane lipid homeostasis under diabetic conditions. Conversely, T2D samples showed a relative increase in fatty acyls and polyketides, indicating a possible shift towards altered fatty acid metabolism and secondary metabolite accumulation. Minor lipid classes such as saccharolipids and sphingolipids were detected at relatively low levels in both groups. To enhance lipid class coverage, we also analyzed the samples in negative ion mode (Figure 7 b). Consistent with the positive mode results, WT samples retained higher levels of glycerophospholipids and glycerolipids. However, the overall lipid signal intensity was lower in the negative mode. Interestingly, T2D samples in negative ion mode exhibited increased levels of fatty acyls and polyketides compared to WT, reinforcing the notion of diabetes induced remodeling of the lipidome. Sterol lipids and prenol lipids were consistently detected in both ionization modes across both genotypes, with no marked differences. Collectively, these results suggest that T2D induces substantial alterations in lipid class distribution, particularly marked by reductions in membrane associated lipids and elevations in lipid classes involved in energy storage and oxidative stress, which may contribute to metabolic dysfunction in the diabetic state.

**Figure 7.**
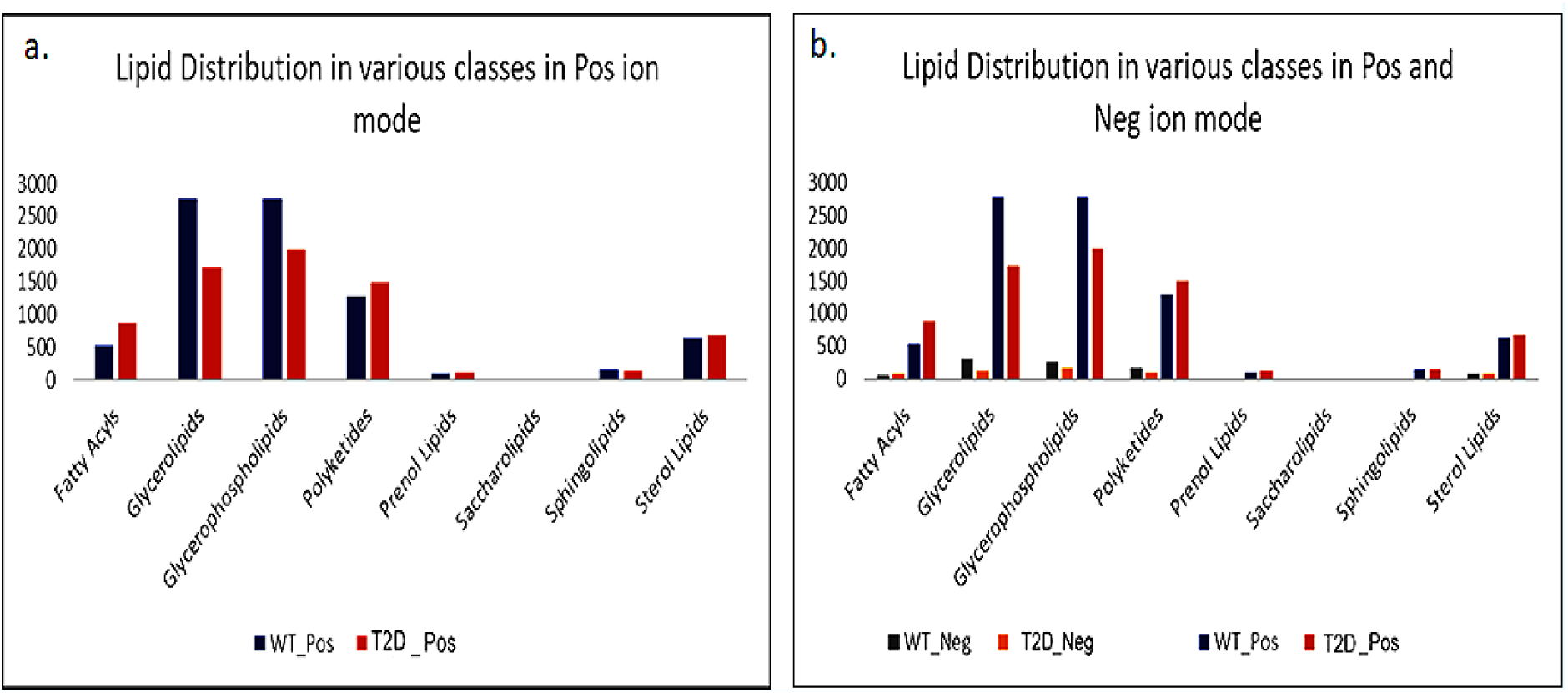
Lipidomic Profiling of WT and T2D Flies: The figure depicts lipid class distribution, with panel (a) representing positive ion mode and panel (b) showing both ion modes.

### 3.10 Random Forest Analysis, Heat Maps, and Biomarker Identification

To identify potential lipid biomarkers associated with T2D, a combination of biomarker analysis, random forest modeling, and receiver operating characteristic (ROC) curve analysis was employed. The biomarker analysis evaluated the sensitivity and specificity of individual lipid species using the area under the ROC curve (AUC) values. The lipid molecules with AUC values exceeding 0.93 and statistically significant log₂ fold change (log₂FC) p-values between WT and T2D adult flies were considered for further analysis. The lipids biomarkers were further filtered based on a stringent differential expression threshold of log₂FC > 1 or < −1, indicating substantial upregulation or downregulation. This approach led to the identification of 32 and 8 free fatty acids in positive (+ESI) and negative (-ESI) ionization modes, respectively (Tables S3 and S4). Random forest analysis and multivariate ROC curve modeling were conducted to refine biomarker selection and assess predictive performance. To further evaluate the diagnostic relevance of identified lipid species in T2D, multivariate receiver operating characteristic (ROC) analysis was conducted in both positive (+ESI) and negative (−ESI) ionization modes (Figure 9). The ROC plots (Figure 8 a & b) demonstrated high classification performance across six models with varying numbers of features. Area under the curve (AUC) values ranged from 0.89 to 1.0, accompanied by narrow confidence intervals (CI), indicating reliable sensitivity and specificity of the selected lipid features in distinguishing T2D from WT flies. The predictive accuracy curves (Figure 8 c & d) revealed a progressive increase in classification accuracy as the number of features increased. Notably, predictive accuracy approached 100% when 20–32 features were used in +ESI mode, and with as few as 5–7 features in −ESI mode, demonstrating high model efficiency even with minimal input. Feature selection frequency analysis using Monte Carlo Cross-Validation (MCCV) is shown in (Figure 8 e & f). Finally, the MCCV prediction score plots (Figure 8 g & h) confirm accurate classification of individual samples into T2D and WT groups based on predicted class probabilities. The clear separation between classes underscores the predictive strength and generalizability of the selected lipid features across both ionization modes. Overall, the integration of multivariate ROC analysis and MCCV provides strong evidence for a core set of lipids with high diagnostic potential. These discriminatory lipid biomarkers, especially those involved in mitochondrial structure and phosphoinositide metabolism, may serve as valuable indicators of metabolic dysregulation in T2D. To identify the most discriminative lipid species associated with T2D in Drosophila, integrated analyses involving receiver operating characteristic (ROC) curves (AUC), t-tests, and fold change were performed.

**Figure 8.**
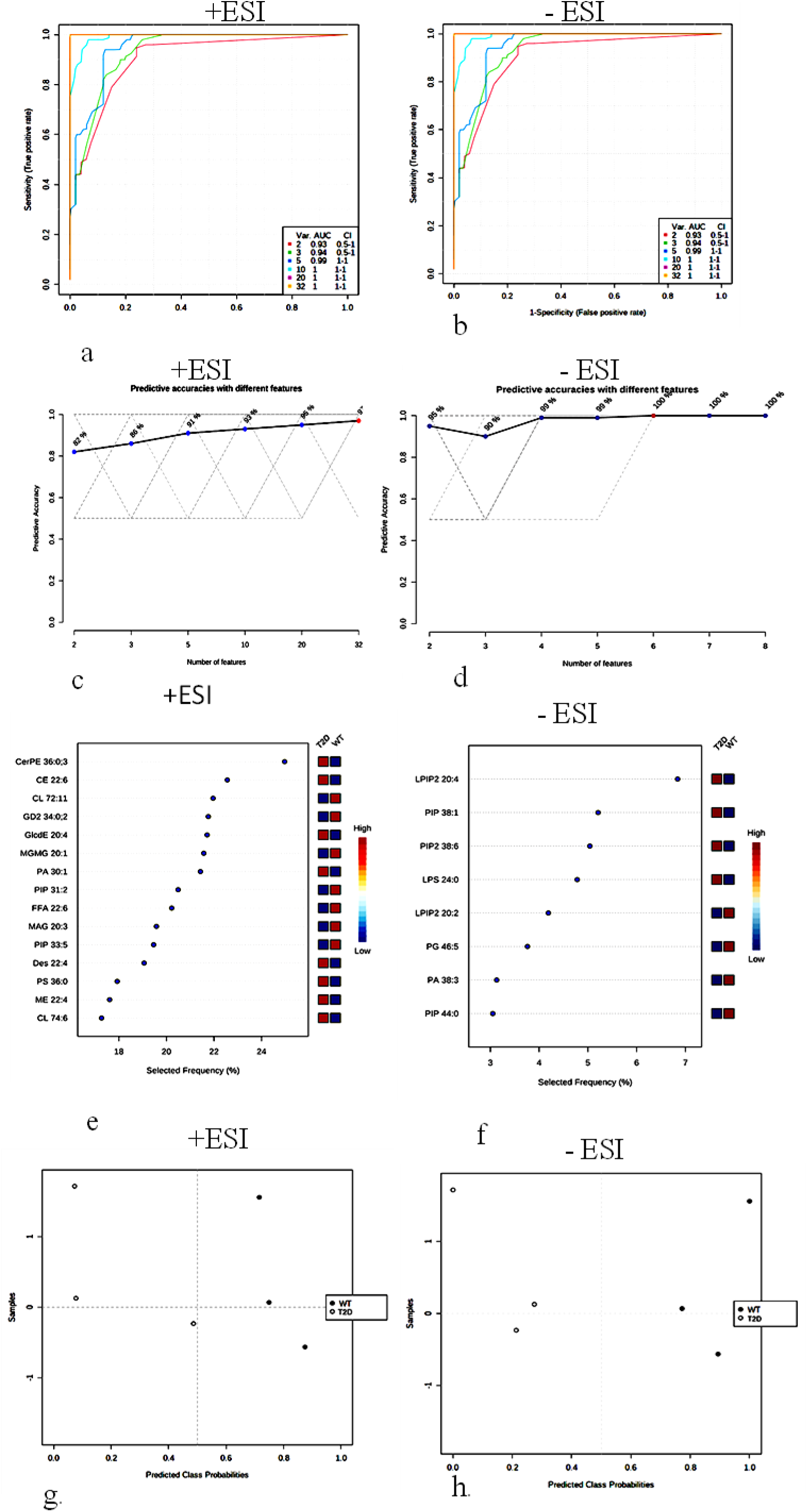
Multivariate receiver operating characteristic (ROC) analysis presented the number of features, AUCs, and confidence intervals across six models (a & b). ROC curves (c & d) illustrated predictive accuracies with varying features. The percentage frequency of selected metabolites was depicted, with a VIP plot highlighting the most discriminating metabolites in order of descending importance (e & f). Monte Carlo cross-validation (MCCV) analysis was employed to predict T2D and WT flies in both +ESI and -ESI ion modes (g & h).

**Figure 9.**
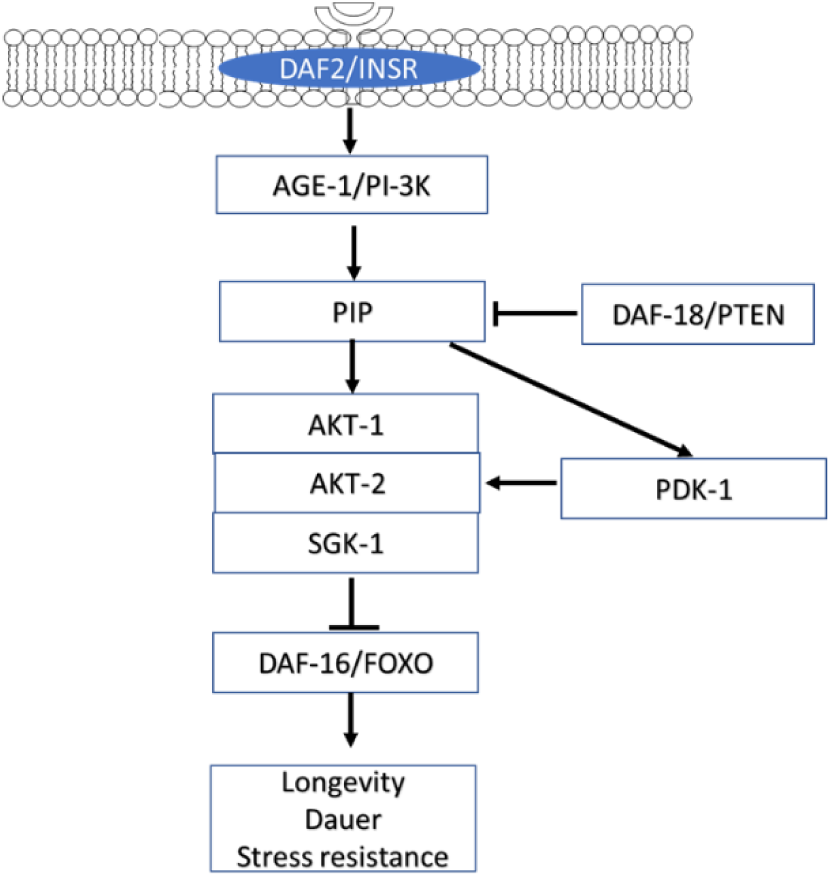
Insulin signaling pathway: Insulin-like peptides (Dilp1-8) bind to DAF-2/DIR receptors, activating β-subunit tyrosine kinase activity and downstream signaling. PI3K converts PIP2 to PIP3, regulated by PTEN. PIP3 recruits PDK1 and Akt1, leading Akt1 phosphorylation, which inhibits DAF-16/FOXO, modulating longevity, dauer formation, and stress resistance. Akt1 is the central kinase in the IIS pathway.

### 3.11 Dysregulation of Insulin Receptor and PI3K Pathway in a *Drosophila* T2D Model: A Lipidomic Perspective

We identified three altered pathways insulin receptor, PI3K, and associated signaling pathways in T2D model [33, 34]. These pathway was compromised in T2D by knockdown of InR. The pathway suggests that the activation of DAF2/INSR leads to a cascade involving AKT-1, AKT-2, and PDK-1, which ultimately influences DAF-16/FOXO. This transcription factor is crucial for enhancing longevity and stress resistance in the organism. The regulation by DAF-18/PTEN indicates a control mechanism that can modulate the pathway activity (Figure 9).

## 4.0 Discussion

In this study, we developed a *Drosophila* model of Type 2 Diabetes (T2D) by using the *dilp2-GAL4>UAS-InR^RNAi^*system to knockdown the insulin receptor (InR) specifically in insulin-producing cells (IPCs) of the brain. Although the present model exhibits several metabolic abnormalities that resemble features of T2D, IPC specific InR knockdown should not be considered direct representation of the pathogenesis of human T2D. In mammals, T2D is a complex systemic disorder characterized primarily by peripheral insulin resistance in tissues such as skeletal muscle, liver, and adipose tissue, accompanied by compensatory and, subsequently, impaired pancreatic β-cell function. In contrast, our Drosophila model specifically reduces InR expression in brain insulin-producing cells (IPCs) and is intended to investigate the consequences of impaired insulin signaling within these cells and its effects on systemic metabolic homeostasis. The *Drosophila* IPCs are functionally analogous to pancreatic β-cells in their production of DILPs, but the organization and regulation of the insulin system differ substantially between flies and mammals [34,35]. Therefore, the metabolic phenotypes observed following IPC specific *InR* knockdown should be interpreted as consequences of impaired insulin signaling in IPCs rather than as a complete recapitulation of human T2D pathogenesis.

Evidence from *Drosophila* indicates that insulin signaling within IPCs can participate in feedback regulation of DILP expression. Broughton demonstrated that manipulation of dInR and dFoxO in the median neurosecretory cells (mNSCs), which include the major DILP producing IPCs, regulates the compensatory increase in *dilp3* expression, supporting an autocrine role for insulin signaling in the regulation of DILP expression [35]. This finding provides a mechanistic basis for feedback regulation of DILP production within IPCs, however, it does not establish that IPC specific *InR* impairment is sufficient to induce diabetes. In the present study, we therefore interpret the observed metabolic abnormalities as consequences of disrupted IPC insulin signaling and altered systemic metabolic homeostasis, rather than as evidence that this model reproduces the complete causal sequence of human T2D [36, 37]. Our model exhibits hallmark T2D like phenotypes, including hyperglycemia increase in glucose, elevated trehalose, triglycerides and increased oxidative stress, consistent with mammalian T2D [38].

A key finding is the reduction in InR mRNA expression in IPCs, coupled with compensatory upregulation of *dilp2*, *dilp3* and *dilp5* reflecting hyperinsulinemia like responses seen in early T2D [35]. This contrasts with insulin deficiency models, such as IPC ablation, which reduce Dilp levels and mimic T1D[34]. The reduced GFP expression in T2D larval brain lobes, driven by *dilp2-GAL4*, indicates IPC dysfunction rather than cell loss, as confirmed by sustained dilp expression. Unexpectedly, we observed a increase in pAkt levels in T2D flies, contrary to the expected decrease in insulin resistance models. This may reflect compensatory activation of the PI3K/Akt pathway due to elevated Dilps or tissue specific signaling in the fly head, as insulin signaling is critical for neuronal function [39]. We quantified total pAkt levels, revealing suggesting that pAkt elevation is driven by increased Dilp signaling. These findings highlight the complexity of insulin signaling in *Drosophila* and underscore the model utility for dissecting T2D like mechanisms.

We observed a increase in H₂O₂ levels, increase in SOD activity, increase in catalase activity, and increase in lipid peroxidation in T2D flies compared to WT. These results indicate heightened ROS production driven by chronic hyperglycemia, consistent with studies in mammalian T2D models where elevated glucose promotes ROS generation, overwhelming antioxidant defenses[40]. The increased H₂O₂ levels in our model suggest mitochondrial dysfunction, likely exacerbated by impaired insulin signaling in IPCs, which disrupts glucose homeostasis and energy metabolism [41]. This aligns with findings by Baenas & Wagner who reported elevated ROS in *Drosophila* T2D like models induced by high sugar diets, linking hyperglycemia to oxidative stress[42]. The marked increase in SOD and catalase activities reflects a compensatory response to neutralize superoxide radicals and hydrogen peroxide, respectively, mitigating oxidative damage in T2D flies. Pejin et al. observed similar upregulation of SOD and catalase in human T2D patients, suggesting that these enzymes are activated to counteract ROS induced stress. However, Dworzański et al. reported elevated SOD and catalase in T2D patients with Epstein-Barr virus infection, consistent with our findings, but noted that such increases may be insufficient to fully restore redox balance[11, 43]. In our model, the elevated LPO indicates that the antioxidant response fails to prevent lipid membrane damage, a key contributor to T2D complications such as β-cell dysfunction and insulin resistance [44]. This is supported by Mandal et al. who found increased MDA levels in T2D patients, correlating with hyperglycemia induced lipid peroxidation. Our lipidomic analysis further complements these findings, as elevated lysophosphatidylserine (LPS 24:0) suggests lipotoxic stress and membrane compromise, consistent with oxidative damage to mitochondrial membranes [45]. The oxidative stress in our model integrates with other T2D phenotypes, amplifying metabolic dysregulation. Hyperglycemia and dyslipidemia (elevated triglycerides) exacerbate ROS production, creating a feedback loop that worsens insulin resistance, as described in mammalian systems [16]. These antioxidant enzymes elevated or steady levels might be a result of their activation in response to oxidative stress and the peroxidation of polyunsaturated fatty acids in cell membranes, which are frequently seen in T2D [41, 43, 46]. Dysregulation of lipid metabolism significantly contributes to the pathogenesis and metabolic control of T2D. Dyslipidemia, characterized by abnormal lipid levels, is frequently observed in T2D patients, contributing to metabolic dysfunction. The recent studies have identified specific lipid molecules that not only indicate the presence of T2D but also serve as potential biomarkers for its treatment of dyslipidemia, the elevated levels of triglycerides and LDL-c are common in T2D patients, while HDL-c levels are often reduced [47]. Research has highlighted significant changes in lysophosphatidylcholine, sphingomyelin, and ceramide levels, which are associated with glucose and lipid metabolism[48, 49]. To elucidate the lipidomic alterations associated with T2D like phenotype, we performed comprehensive lipid profiling in *Drosophila* using both positive and negative electrospray ionization (ESI) modes. As shown in Figure 4 a, major lipid classes including glycerolipids, glycerophospholipids, and polyketides were markedly altered in T2D flies compared to WT in positive ion mode. This shift was further validated across both ion modes (Figure 4 b), where glycerophospholipids and polyketides were significantly downregulated in T2D, highlighting the disruption in membrane lipid composition and metabolic flux. We analyzed the insulin signaling cascade (Figure 5), where INSR/DAF-2 signaling modulates downstream targets like PI3K, AKT, and FOXO. Disruption of this pathway in T2D flies may lead to altered lipid metabolism, oxidative stress response, and impaired mitochondrial function hallmarks of metabolic syndrome. The multivariate statistical analyses using PCA and PLS-DA (Figures 6 and 7) revealed a distinct separation between WT and T2D groups in both ionization modes, confirming global lipidomic divergence. The scree plots and score plots (Figures 4 a–6 f) highlight principal components for a majority of the variance, indicating robust clustering. The 3D score and loading plots further confirmed that the discriminating features correspond to lipid molecules with differential abundance.

The volcano plots (Figures 5 a–b) and heatmaps (Figures 5 c–d) provided visual affirmation of statistically significant and biologically relevant lipid species with consistent patterns across replicates. Notably, a subset of lipids including PG 34:0, PA 38:3, and PIP 38:6 showed substantial fold changes and clear clustering, highlighting them as potential biomarkers. Receiver Operating Characteristic (ROC) analyses (Figures 8 a–b) demonstrated that several lipid features had high predictive power (AUC = 1.0), and feature selection plots (Figures 8 c–d) confirmed that even a limited number of lipids could discriminate between T2D and WT with high accuracy. The frequency plots (Figures 8 g–h) reinforced the biomarker potential of individual lipids. Predicted class probabilities (Figures 8 i–j) further demonstrated the high classification confidence between T2D and WT groups. Finally, targeted ROC analyses (Figure 6) for top discriminating lipids reaffirmed their diagnostic relevance. Lipids such as PG 34:0, PG 36:4, PA 38:3, PIP 38:1, and LPS 24:0 showed perfect separation between the groups with an AUC of 1.0, suggesting their potential use as metabolic indicators of insulin resistance and lipid dysregulation in diabetes. Furthermore, VIP score plots (Figures 8 d, 8 h) identified key discriminatory lipids such as phosphatidylinositol phosphates (PIPs), lysophosphatidylserines (LPS), and phosphatidic acids (PAs), which were among the top contributors to group separation. Together, these data strongly indicate that the diabetic condition in *Drosophila* significantly alters the lipidome through PI3K/AKT pathway perturbation, contributing to cellular stress and metabolic imbalance. These lipidomic biomarkers may offer novel insights into disease progression and therapeutic targets for T2D management. Two distinct lipid profiles have been identified; one associated with lower T2D risk (including lysophospholipids and sphingomyelins) and another linked to higher risk composed of triacylglycerols and diacylglycerols [48–51]. These discriminatory lipid species likely include mitochondrial-associated phospholipids such as phosphatidylglycerols (PGs), phosphatidic acid (PA), phosphoinositides (PIPs), and lysophosphatidylserine (LPS), all of which play critical roles in membrane integrity, energy metabolism, and insulin signaling. The consistent pattern across both ion modes confirms that T2D induces widespread and distinct lipidomic remodeling, potentially contributing to mitochondrial dysfunction and metabolic dysregulation.

In a comprehensive metabolomics investigation, Peddinti, G. discerned that [Hyp3]-BK, α-tocopherol, and X-13435 exhibited negative correlation with the incidence of T2D, whereas glucose, mannose, α-HB, and X-12063 displayed of positive correlation with the same condition[52]. Suvitaival, T. observed that among those who were moving towards T2D, there were higher concentrations of triacylglycerols and diacylphospholipids and lower levels of alkyl acyl phosphatidylcholines. Specifically, triacylglycerol TG(17:1/18:1/18:2), phosphatidylcholines PC(32:1), PC(34:2e), and PC(36:1), as well as lysophosphatidylcholine acyl C18:2 (LysoPC(18:2)), were incorporated into the comprehensive model alongside metabolic risk variables to predict the progression of T2D [53]. The predictive significance of indicators showing a negative correlation with illness risk is being emphasised for the first time in this study. Contrary to this findings, previous studies showed that mannose was more strongly linked to an elevated risk of T2D than glucose. Zhang et.a., reported the quantitative analysis of plasma samples to measure the concentrations of eight lipid metabolites: palmitic acid, stearic acid, arachidonic acid, PC (18:2/18:0), PC (20:4/18:2), PC (18:2/18:2), SM(d18:0/18:1), and LysoPC (18:0/0:0). The discriminatory effectiveness each biomarker was assessed using ROC curve analysis, with performance quantified by AUC [18]. Ma et al. reported eleven free fatty acids (FFAs) within an OPLS-DA model, highlighting their significant contribution to the differentiation of metabolic profiles across experimental groups. The analysis revealed that nine FFAs C14:0, C18:1, C20:1, C18:2, C20:2, C20:3, C18:3, C20:5, and C22:6 exhibited statistically significant differences (P < 0.05) between WT and T2D[54]. Our lipidomic analysis identified significant dysregulation of six mitochondrial associated phospholipids in T2D like models, highlighting mechanisms of mitochondrial dysfunction. Elevated phosphatidylglycerol (PG 34:0 and PG 34:4) suggests disturbed cardiolipin biosynthesis PG is a precursor for cardiolipin, which maintains inner mitochondrial membrane integrity and supports oxidative phosphorylation through electron transport chain (ETC) stabilisation[55]. Increased phosphatidic acid (PA 38:3) points to enhanced mitochondrial fission and mitophagy, consistent with stress responses observed in insulin resistance via altered mitochondrial dynamics[56]. The reduction in PIP 38:1 and PIP2 38:6 reflects impaired PI3K/Akt signaling and dysfunctional calcium handling at mitochondrial associated membranes both central to energy metabolism and disrupted in T2D. Finally, the decrease in lysophosphatidylserine (LPS 24:0) likely indicates lipid peroxidation, membrane compromise, and initiation of apoptotic signaling factors that accentuate mitochondrial failure, as seen in β-cell demise and diabetic pathophysiology. Together, these lipid alterations underscore a coordinated remodeling of mitochondrial membranes and signaling networks, pinpointing specific lipid species as both biomarkers and mechanistic drivers of mitochondrial dysfunction in T2D[45, 57]. The PA 38:3, a precursor to other harmful lipid molecules, can also contribute to insulin resistance. Moreover, advancing our knowledge of this widespread metabolic disorder provides strong support for our findings[58]. Phospholipids and lipopolysaccharides (LPS) are intricate lipid compounds linked to metabolic health and inflammation, crucial factors in developing T2D. PIP38:1 is a phosphatidylinositol phospholipid found in cell membranes and is critical for insulin signaling. PIP2 38:6, phosphatidylinositol bisphosphate, influences insulin receptor activation and signaling, shedding light on its role in insulin sensitivity and glucose regulation [59, 60].

In this study, we demonstrate the utility of metabolomics in elucidating the mechanisms by which T2D impacts critical pathways. Despite its global prevalence, the precise pathophysiology of T2D remains poorly understood. Using a systematic UPLC/MS based metabolomics approach, we uncovered significant insights into T2D. These results highlight the effectiveness of metabolomics in identifying T2D biomarkers. The non invasive, real time analysis of lipid metabolites in a T2D fly model offers valuable insights into disease mechanisms, supporting biomarker discovery and the development of targeted therapies.

## 5.0 Conclusion

In this study, we established a novel *Drosophila* T2D like model by specifically knockdown the *InR* in insulin-producing cells (IPCs) of the brain. This genetic perturbation induced a range of T2D-like phenotypes, including hyperglycemia, dysregulated lipid profiles, increased oxidative stress, and altered expression of insulin signaling components. Using a UPLC-ESI-MS-based untargeted lipidomics approach, we identified distinct alterations in multiple lipid classes, with six phosphatidylglycerol species emerging as potential biomarkers associated with the T2D state. These findings demonstrate the utility of *Drosophila* as a powerful model to uncover metabolic signatures of T2D and provide new insights into lipid dysregulation in insulin resistant conditions. While further validation in mammalian systems is warranted, our study lays the groundwork for exploring evolutionarily conserved lipid biomarkers and pathways implicated in T2D pathogenesis.

## Credit authorship contribution statement

Prabhat Kumar: Conceptualization, Investigation, Visualization, Validation, Writing – original draft; Pradeep Kumar: Sample preparation, Data acquisition, Data analysis: Rohit Kumar; Sample preparation, Data acquisition, Data analysis; Brijesh Singh Chauhan: Data analysis, Writing – review and editing; Zeeshan Fatima: UPLC-MS facility, Data acquisition, Data analysis, Computational analysis, Writing – review and editing; Saripella Srikrishna: Conceptualization, Funding acquisition, Resources, Supervision, Writing – review and editing.

## Acknowledgments

The authors highly acknowledge Dr. Jishy Varghese for providing dilp2-Gal4 stock and BDSC. ISLS-BHU for providing a confocal facility. P K dramatically acknowledges the financial support from the CSIR-JRF-SRF, IoE, BHU Sathi.

## Abbreviation

T2D: Type 2 diabetes
InR: insulin receptor
Dilp: *Drosophila* insulin-like peptide
SOD: Sodium oxide dismutase
TBARS: Thiobarbituric acid reactive substances
FFA: Free Fatty Acid
PIP: Phosphatidylinositol phosphate
PG: Phosphatidylglycerol
PA: Phosphatidic acid
LPS: Lipopolysaccharides
TAG: Triacylglycerols
+ESI: Positive Electrospray Ionization
−ESI: Negative Electrospray Ionization
UPLC-QTOF-MS: Ultra-performance liquid chromatography coupled with quadrupole time-of-flight mass spectrometry
PCA: Principal Component Analysis
PLS-DA: Partial Least Squares Discriminant Analysis
VIP: Scores Variable Importance in Projection Scores

