## Supplementary material for "UPLC-ESI-MS based lipidomics revealed novel biomarkers in insulin receptor knockdown induced type 2 diabetes model of *Drosophila*": Supplimentary Information.docx

**Table: S 1** The distribution of identified lipids across different classes was determined using MS-LAMP.

| S.No Class | | WT (Positive) | T2D (Positive) |
| --- | --- | --- | --- |
| 1 | Fatty acyls | 537 | 872 |
| 2 | Glycerolipids | 2778 | 1723 |
| 3 | Glycerophospholipids | 2782 | 1997 |
| 4 | Polyketides | 1287 | 1503 |
| 5 | Prenol lipids | 107 | 121 |
| 6 | Saccharolipids | 7 | 2 |
| 7 | Sphingolipids | 159 | 145 |
| 8 | Sterol lipids | 639 | 688 |
| Class | | **WT (Negative)** | **T2D (Negative)** |
| 1 | Fatty acyls | 67 | 73 |
| 2 | Glycerolipids | 310 | 137 |
| 3 | Glycerophospholipids | 277 | 172 |
| 4 | Polyketides | 184 | 111 |
| 5 | Prenol lipids | 11 | 5 |
| 6 | Saccharolipids | 0 | 0 |
| 7 | Sphingolipids | 24 | 6 |
| 8 | Sterol lipids | 82 | 77 |

**Table S 2.** The concentration of free fatty acids, detected using positive ion modes by Ultra-High-Pressure Liquid Chromatography-Mass Spectrometry (UPLC-MS), exhibited a significant difference. The data were subjected to cluster analysis for univariate biomarker evaluation, incorporating Area Under the Curve (AUC), log2 fold change, K-means clustering, and independent t-tests, comparing WT and T2D flies.

| S.No. | Free fatty acids | The area under the curve | t-test | Log2 fold change | Cluster |
| --- | --- | --- | --- | --- | --- |
| 1 | CL 60:3 | 1.00 | 0.03 | 1.51 | 18.00 |
| 2 | PS 36:0 | 1.00 | 0.07 | -1.62 | 23.00 |
| 3 | PS 36:2 | 1.00 | 0.07 | -1.51 | 9.00 |
| 4 | PS 36:3 | 1.00 | 0.02 | -1.88 | 30.00 |
| 5 | DGPP 38:4 | 1.00 | 0.06 | -1.19 | 20.00 |
| 6 | DGPP 36:6 | 1.00 | 0.08 | 1.17 | 4.00 |
| 7 | CL 74:12 | 1.00 | 0.07 | 1.30 | 2.00 |
| 8 | DGPP 38:1 | 1.00 | 0.00 | 2.42 | 11.00 |
| 9 | GlcdE 20:1 | 1.00 | 0.00 | -2.38 | 25.00 |
| 10 | Glcde 22:0 | 1.00 | 0.00 | -2.45 | 27.00 |
| 11 | Glcde 22:4 | 1.00 | 0.05 | -1.46 | 22.00 |
| 12 | MAG 18:0 | 1.00 | 0.08 | -1.13 | 7.00 |
| 13 | ME 22:4 | 1.00 | 0.06 | 1.36 | 16.00 |
| 14 | MMPE 34:4 | 1.00 | 0.01 | -2.62 | 17.00 |
| 15 | PG 30:2 | 1.00 | 0.05 | -0.75 | 13.00 |
| 16 | PG 34:0 | 1.00 | 0.00 | 2.33 | 31.00 |
| 17 | PG 34:4 | 1.00 | 0.00 | 2.35 | 5.00 |
| 18 | CL 74:6 | 1.00 | 0.06 | 1.34 | 29.00 |
| 19 | PIP 30:3 | 1.00 | 0.07 | 1.23 | 1.00 |
| 20 | PIP 38:1 | 1.00 | 0.01 | 3.22 | 6.00 |
| 21 | Des 22:4 | 0.89 | 0.08 | 1.20 | 21.00 |
| 22 | MAG 20:3 | 0.89 | 0.12 | 0.27 | 24.00 |
| 23 | PIP 31:2 | 0.89 | 0.09 | 1.22 | 14.00 |
| 24 | CE 22:6 | 0.78 | 017 | 1.35 | 12.00 |
| 25 | CL 72:11 | 0.78 | 0.11 | -1.09 | 10.00 |
| 26 | FFA 22:6 | 0.78 | 0.11 | -1.07 | 15.00 |
| 27 | GD2 34:0:2 | 0.78 | 0.20 | -0.94 | 3.00 |
| 28 | GlcdE 20:4 | 0.78 | 0.11 | -1.13 | 19.00 |
| 29 | MGMG 20:1 | 0.78 | 0.10 | -1.10 | 10.00 |
| 30 | PA 30:1 | 0.78 | 0.16 | -1.09 | 28.00 |
| 31 | PIP 33:5 | 0.78 | 0.14 | -1.14 | 26.00 |
| 32 | CerPE 36:0:3 | 0.67 | 0.49 | -0.11 | 8.00 |

**Table S 3.** The concentration of free fatty acids, identified using negative ion modes via Ultra-High-Pressure Liquid Chromatography-Mass Spectrometry (UPLC-MS), showed a significant difference. The data were analyzed through cluster methods for univariate biomarker assessment, utilizing Area Under the Curve (AUC), log2 fold change, K-means clustering, and independent t-tests to compare WT and T2D flies.

| S.no | Free fatty acids | The area under the curve | t-test | Log2 fold change | Cluster |
| --- | --- | --- | --- | --- | --- |
| 1 | PA 38:3 | 1.00 | 0.00 | 2.73 | 4.00 |
| 2 | PIP 44:0 | 1.00 | 0.00 | 2.49 | 4.00 |
| 3 | LPIP 20:2 | 1.00 | 0.03 | 0.56 | 1.00 |
| 4 | PIP 38:1 | 1.00 | 0.04 | -2.55 | 2.00 |
| 5 | PIP2 38:6 | 1.00 | 0.05 | -2.42 | 2.00 |
| 6 | LPS 24:0 | 1.00 | 0.05 | -2.44 | 2.00 |
| 7 | PG 46:5 | 1.00 | 0.01 | 0.98 | 3.00 |
| 8 | LPIP2 20:4 | 0.67 | 0.53 | -1.66 | 5.00 |

**Table S 4.** A significant difference in potential biomarkers was detected using positive ion modes via Ultra-High-Pressure Liquid Chromatography-Mass Spectrometry (UPLC-MS). The potential biomarkers were identified based on m/z values obtained through t-tests, fold change analysis, and the absence of lipid indicated by a ‘0’ intensity. Lipids were identified using both MS-LAMP (with a WR threshold of 0.01) and the Lipid Maps database.

| S.No. | Mass | Lipid | WT | WT | WT | T2D | T2D | T2D |
| --- | --- | --- | --- | --- | --- | --- | --- | --- |
| 1 | 751.6 | PG 34:0 | 0 | 0 | 0 | 1720291.5 | 1709556.1 | 1704035.9 |
| 2 | 743.3 | PG 34:4 | 0 | 0 | 0 | 1873070.3 | 1880734.4 | 1885650.2 |

**Table S 5.** A significant difference in potential biomarkers was detected using negative ion modes through Ultra-High-Pressure Liquid Chromatography-Mass Spectrometry (UPLC-MS). The potential biomarkers were identified based on m/z values obtained from t-tests and fold change analysis, with a ‘0’ intensity indicating the absence of lipids. Lipids were identified using both MS-LAMP (with a WR threshold of 0.01) and the Lipid Maps database.

|  | Mass | Lipid | WT | WT | WT | T2D | T2D | T2D |
| --- | --- | --- | --- | --- | --- | --- | --- | --- |
| 1 | 725.5 | PA 38:3 | 0 | 0 | 0 | 98923.5 | 139853.4 | 154706.8 |
| 2 | 485.3 | PIP 38:1 | 186308.2 | 203448.3 | 155337.9 | 0 | 0 | 0 |
| 3 | 520.2 | PIP2 38:6 | 295766.6 | 306321.8 | 273425.5 | 0 | 0 | 0 |
| 4 | 608.4 | LPS 24:0 | 303013 | 281992.3 | 258256.5 | 0 | 0 | 0 |

**Table S 6.** Biomarkers were identified through the analysis of lipid molecules using Area Under the Curve (AUC), t-tests, and fold change assessments.

| PG 34:0 | AUC 1.0 | | T-Test 4.1092E-5 | | | | | Log 2FC 2.3281 | | | | | | |
| --- | --- | --- | --- | --- | --- | --- | --- | --- | --- | --- | --- | --- | --- | --- |
| cut-off value | **sensitivity** | | **specificity** | | | | **Sum of (sens. + spec.) likelihood ratios (LR+ and LR-)** | | | | | | | |
| Infinity | 1.0 | | 0.0 | | | | 1.0 | | | 1.0 | | | NaN | |
| 1.0191 | 1.0 | | 0.333333 | | | | 1.33333 | | | 1.5 | | | 0.0 | |
| 0.834483 | 1.0 | | 0.666666 | | | | 1.66667 | | | 3.0 | | | 0.0 | |
| -0.0975451 | 1.0 | | 1.0 | | | | 2.0 | | | Infinity | | | 0.0 | |
| -0.89132 | 0.666667 | | 1.0 | | | | 1.66667 | | | Infinity | | | 0.333333 | |
| -0.921553 | 0.333333 | | 1.0 | | | | 1.33333 | | | Infinity | | | 0.6666667 | |
| PG 34:4 | **AUC 1.0** | | **T-Test 9.8817E-5** | | | | **Log 2FC. 2.3459** | | | | | | | |
| cut-off value | **sensitivity** | **specificity** | | | **The sum of (sens. + spec.)** | | | | | | **likelihood ratios (LR+ and LR-)** | | | |
| -Infinity | 1.0 | 0.0 | | | 1.0 | | | | | | 1.0 NaN | | | |
| 1.0191 | 1.0 | 0.666667 | | | 1.66667 | | | | | | 1.5 | | | 0.0 |
| 0.834483 | 1.0 | 0.333333 | | | 1.33333 | | | | | | 3 | | | 0.0 |
| -0.0975451 | 1.0 | 1.0 | | | 2.0 | | | | | | Infinity | | | 0.0 |
| -0.89132 | 0.666667 | 1.0 | | | 1.66667 | | | | | | Infinity | | | 0.666667 |
| -0.921553 | 0.333333 | 1.0 | | | 1.33333 | | | | | | Infinity | | | 0.333333 |
| -Infinity | 0.0 | 1.0 | | | 1.0 | | | | | | NaN | | | 1.0 |
| PA 38:3 | **AUC 1.0** | | | **T-Test 0.003301** | | | | | | | **Log 2FC. 2.7289** | | | |
| cut-off value | **sensitivity** | | | **specificity** | | **The sum of (sens. + spec.)** | | | **likelihood ratios (LR+ and LR-)** | | | | | |
| Infinity | 1.0 | | | 0.0 | | 1.0 | | | 1.0 | | | NaN | | |
| 1.13615 | 1.0 | | | 0.666667 | | 1.66667 | | | 1.5 | | | 0.0 | | |
| 0.669737 | 1.0 | | | 0.333333 | | 1.33333 | | | 3 | | | 0.0 | | |
| -0.263885 | 1.0 | | | 1.0 | | 2.0 | | | Infinity | | | 0.0 | | |
| -0.866281 | 0.666667 | | | 1.0 | | 1.66667 | | | Infinity | | | 0.666667 | | |
| -0.872266 | 0.333333 | | | 1.0 | | 1.33333 | | | Infinity | | | 0.333333 | | |
| -Infinity | 0.0 | | | 1.0 | | 1.0 | | | NaN | | | 1.0 | | |
| PIP 38:1 | **AUC 1.0** | | | **T-Test 0.042802** | | | | | | | **Log 2FC. -2.5481** | | | |
| cut-off value | **sensitivity** | | | **specificity** | | **The sum of (sens. + spec.)** | | | **likelihood ratios (LR+ and LR-)** | | | | | |
| -Infinity | 1.0 | | | 0.0 | | 1.0 | | | 1.0 | | | NaN | | |
| -1.07974 | 1.0 | | | 0.333333 | | 1.33333 | | | 1.5 | | | 0.0 | | |
| -0.434123 | 1.0 | | | 0.666667 | | 1.666667 | | | 3 | | | 0.0 | | |
| -0.0248287 | 1.0 | | | 1.0 | | 2.0 | | | Infinity | | | 0.0 | | |
| 0.531668 | 0.666667 | | | 1.0 | | 1.66667 | | | Infinity | | | 0.666667 | | |
| 1.10457 | 0.333333 | | | 1.0 | | 1.33333 | | | Infinity | | | 0.333333 | | |
| Infinity | 0.0 | | | 1.0 | | 1.0 | | | NaN | | | 1.0 | | |
| PIP2 38:6 | **AUC 1.0** | | | **T-Test 0.046065** | | | | | | | **Log 2FC. -2.4159** | | | |
| cut-off value | **sensitivity** | | | **specificity** | | **The sum of (sens. + spec.)** | | | **likelihood ratios (LR+ and LR-)** | | | | | |
| Infinity | 1.0 | | | 0.0 | | 1.0 | | | 1.0 | | | NaN | | |
| -1.14961 | 1.0 | | | 0.333333 | | 1.33333 | | | 1.5 | | | 0.0 | | |
| -0.353344 | 1.0 | | | 0.666667 | | 1.666667 | | | 3 | | | 0.0 | | |
| 0.151456 | 1.0 | | | 1.0 | | 2.0 | | | Infinity | | | 0.0 | | |
| 0.621582 | 0.666667 | | | 1.0 | | 1.66667 | | | Infinity | | | 0.333333 | | |
| 0.998157 | 0.333333 | | | 1.0 | | 1.33333 | | | Infinity | | | 0.666667 | | |
| Infinity | 0.0 | | | 1.0 | | 1.0 | | | NaN | | | 1.0 | | |
| LPS 24:0 | **AUC 1.0** | | | **T-Test 0.047748** | | | | | | | **Log 2FC. -2.4441** | | | |
| cut-off value | **sensitivity** | | | **specificity** | | **The sum of (sens. + spec.)** | | | **likelihood ratios (LR+ and LR-)** | | | | | |
| Infinity | 1.0 | | | 0.0 | | 1.0 | | | 1.0 | | | NaN | | |
| -1.12507 | 1.0 | | | 0.333333 | | 1.33333 | | | 1.5 | | | 0.0 | | |
| -0.371321 | 1.0 | | | 0.666667 | | 1.666667 | | | 3 | | | 0.0 | | |
| 0.106526 | 1.0 | | | 1.0 | | 2.0 | | | Infinity | | | 0.0 | | |
| 0.504388 | 0.666667 | | | 1.0 | | 1.66667 | | | Infinity | | | 0.333333 | | |
| 1.01855 | 0.333333 | | | 1.0 | | 1.33333 | | | Infinity | | | 0.666667 | | |
| Infinity | 0.0 | | | 1.0 | | 1.0 | | | NaN | | | 1.0 | | |

**Table S 7.** Paired fold change analysis was conducted in both positive and negative ion modes.

|  | Positive ions modes | | | Negative ions modes | | |
| --- | --- | --- | --- | --- | --- | --- |
| S.No. | **Peaks mz/rt** | **Fold change** | **Log2 FC** | **Peaks mz/rt** | **Fold change** | **Log2 FC** |
| 1 | 487.3 | 8.3493 | 3.0617 | 725.5 | 9.7594 | 3.2868 |
| 2 | 726.5 | 0.14829 | -2.7535 | 528.35 | 8.3923 | 3.0691 |
| 3 | 397.35 | 0.16605 | -2.5909 | 1013.35 | 7.8438 | 2.9716 |
| 4 | 367.35 | 0.17457 | -2.5181 | 735.4 | 4.9769 | 2.3152 |
| 5 | 811.5 | 4.8491 | 2.2777 | 485.3 | 0.25858 | -1.9513 |
| 6 | 852.45 | 4.7652 | 2.2525 | 496.2 | 0.27156 | -1.8806 |
| 7 | 839.5 | 4.7283 | 2.2413 | 904.55 | 0.27378 | -1.8689 |
| 8 | 743.5 | 4.6175 | 2.2071 | 608.4 | 0.27719 | -1.851 |
| 9 | 751.6 | 4.5614 | 2.1895 | 520.2 | 0.28376 | -1.8172 |
| 10 | 896.5 | 0.24556 | -2.0259 | 1121.4 | 3.3973 | 1.7644 |
| 11 | 786.55 | 0.2486 | -2.0081 | 1127.5 | 3.3403 | 1.74 |
| 12 | 899.4 | 0.25215 | -1.9876 | 1109.6 | 3.0407 | 1.6044 |
| 13 | 792.55 | 0.29793 | -1.7469 | 1191.7 | 2.9848 | 1.5777 |
| 14 | 348.35 | 0.31755 | -1.6549 | 907.65 | 2.6428 | 1.4021 |
| 15 | 788.55 | 0.31997 | -1.644 | 392.1 | 0.39388 | -1.3442 |
| 16 | 389.3 | 0.33213 | -1.5902 | 479.3 | 2.3818 | 1.252 |
| 17 | 646.45 | 2.6071 | 1.3825 | 391.15 | 2.3464 | 1.2305 |
| 18 | 587.4 | 0.39321 | -1.3466 | 389.05 | 0.44766 | -1.1595 |
| 19 | 805.5 | 0.39977 | -1.3228 |  |  |  |
| 20 | 895.5 | 0.40723 | -1.2961 |  |  |  |
| 21 | 361.35 | 0.41767 | -1.2596 |  |  |  |
| 22 | 376.35 | 0.41829 | -1.2574 |  |  |  |
| 23 | 863.5 | 0.42181 | -1.2453 |  |  |  |
| 24 | 843.45 | 2.3523 | 1.2341 |  |  |  |
| 25 | 951.35 | 0.42639 | -1.2298 |  |  |  |
| 26 | 581.4 | 0.42695 | -1.2279 |  |  |  |
| 27 | 619.45 | 0.42838 | -1.2231 |  |  |  |
| 28 | 714.6 | 2.3338 | 1.2227 |  |  |  |
| 29 | 913.4 | 0.42886 | -1.2214 |  |  |  |
| 30 | 722.45 | 0.43053 | -1.2158 |  |  |  |
| 31 | 347.35 | 2.3085 | 1.207 |  |  |  |
| 32 | 329.2 | 0.43441 | -1.2029 |  |  |  |
| 33 | 741.5 | 2.2705 | 1.183 |  |  |  |
| 34 | 735.55 | 2.2088 | 1.1433 |  |  |  |
| 35 | 429.2 | 2.1453 | 1.1011 |  |  |  |
| 36 | 750.5 | 2.1279 | 1.0894 |  |  |  |
| 37 | 977.35 | 2.1279 | 1.894 |  |  |  |
| 38 | 716.6 | 2.1074 | 1.0755 |  |  |  |
| 39 | 647.6 | 2.1007 | 1.0709 |  |  |  |
| 40 | 825.5 | 0.47652 | -1.0694 |  |  |  |
| 41 | 437.25 | 2.0899 | 1.0635 |  |  |  |
| 42 | 923.45 | 0.48324 | -1.0492 |  |  |  |
| 43 | 801.5 | 2.0598 | 1.0425 |  |  |  |
| 44 | 636.4 | 2.0198 | 1.0142 |  |  |  |

**Table S 8** The significant m/z values were identified through fold change analysis using t-tests in positive ion mode**.**

| S.no. | Peaks mz/rt | t-test | p-value | -log10(p) | FDR |
| --- | --- | --- | --- | --- | --- |
| 1 | 743.5 | 105.15 | 4.9054e-08 | 7.3093 | 8.4373e-06 |
| 2 | 751.6 | 74.45 | 1.9506e-07 | 6.7098 | 1.6775e-05 |
| 3 | 839.5 | 47.509 | 1.1743e-06 | 5.9302 | 6.7326e-05 |
| 4 | 892.45 | 38.902 | 2.6082e06 | 5.5837 | 0.00011215 |
| 5 | 367.35 | -27.694 | 1.0112e-05 | 4.9952 | 0.00034785 |
| 6 | 811.5 | 18.738 | 4.7758e-05 | 4.321 | 0.0013691 |
| 7 | 397.35 | -17.708 | 5.9747e-05 | 4.2237 | 0.0014681 |
| 8 | 726.5 | -6.8267 | 0.0024077 | 2.6184 | 0.051766 |
| 9 | 385.3 | -5.9274 | 0.0040596 | 2.3915 | 0.077584 |
| 10 | 899.4 | -4.4806 | 0.010985 | 1.9592 | 0.18894 |
| 11 | 896.5 | -4.2674 | 0.012977 | 1.8868 | 0.20291 |
| 12 | 786.55 | -4.0831 | 0.01506 | 1.8222 | 0.21586 |
| 13 | 487.3 | 3.9548 | 0.016751 | 1.776 | 0.22162 |
| 14 | 348.35 | -3.6466 | 0.021836 | 1.6608 | 0.26827 |
| 15 | 892.4 | -3.3339 | 0.029 | 1.5376 | 0.33253 |
| 16 | 389.3 | -2.9557 | 0.041732 | 1.3795 | 0.42011 |
| 17 | 646.45 | 2.7956 | 0.049032 | 1.3095 | 0.42011 |
| 18 | 788.55 | -2.6756 | 0.055483 | 1.2558 | 0.42011 |
| 19 | 691.5 | -2.6728 | 0.055644 | 1.2546 | 0.42011 |
| 20 | 880.45 | -2.5861 | 0.060936 | 1.2151 | 0.42011 |
| 21 | 805.5 | -2.5662 | 0.062228 | 1.206 | 0.42011 |
| 22 | 895.5 | -2.4384 | 0.071335 | 1.1467 | 0.42011 |
| 23 | 868.4 | -2.4016 | 0.07423 | 1.1294 | 0.42011 |
| 24 | 792.55 | -2.3736 | 0.076521 | 1.1162 | 0.42011 |
| 25 | 843.45 | 2.2896 | 0.083898 | 1.0762 | 0.42011 |
| 26 | 376.35 | -2.2853 | 0.084292 | 1.0742 | 0.42011 |
| 27 | 951.35 | -2.2809 | 0.084704 | 1.0721 | 0.42011 |
| 28 | 347.35 | 2.249 | 0.087745 | 1.0568 | 0.42011 |
| 29 | 365.25 | -2.2316 | 0.089454 | 1.0484 | 0.42011 |
| 30 | 398.35 | 2.2084 | 0.091789 | 1.0372 | 0.42011 |
| 31 | 741.5 | 2.1722 | 0.095579 | 1.0196 | 0.42011 |
| 32 | 707.55 | -2.1661 | 0.096235 | 1.0167 | 0.42011 |
| 33 | 913.4 | -2.1641 | 0.096443 | 1.0157 | 0.42011 |
| 34 | 619.45 | -2.1371 | 0.099414 | 1.0026 | 0.42011 |

**Table S 9.** The Key m/z values were identified through fold change analysis using t-tests in negative ion mode.

| S.No. | Peaks mz/rt | t-test | p-value | -log10(p) | FDR |
| --- | --- | --- | --- | --- | --- |
| 1 | 725.5 | 26.968 | 1.124e-05 | 4.9492 | 0.00044959 |
| 2 | 904.55 | -21.758 | 2.6398e-05 | 4.5784 | 0.00052796 |
| 3 | 528.35 | 15.046 | 0.00011372 | 3.9442 | 0.0015163 |
| 4 | 1013.35 | 8.9248 | 0.00087149 | 3.0597 | 0.0087149 |
| 5 | 520.2 | -8.3822 | 0.0011081 | 2.9554 | 0.008865 |
| 6 | 496.2 | -6.7895 | 0.0024573 | 2.6095 | 0.014836 |
| 7 | 485.3 | -6.5736 | 0.0029671 | 2.5573 | 0.014836 |
| 8 | 608.4 | -6.454 | 0.0027716 | 2.5277 | 0.014836 |
| 9 | 1127.5 | 3.307 | 0.029739 | 1.5267 | 0.13217 |
| 10 | 1121.4 | 3.1474 | 0.034602 | 1.4609 | 0.13841 |
| 11 | 1191.7 | 2.9592 | 0.041587 | 1.381 | 0.14892 |
| 12 | 1109.6 | 2.8874 | 0.044677 | 1.3499 | 0.14892 |
| 13 | 404.05 | 2.7209 | 0.05294 | 1.2762 | 0.15149 |
| 14 | 735.4 | 2.7194 | 0.053022 | 1.2755 | 0.15149 |
| 15 | 392.1 | -2.4524 | 0.070261 | 1.1533 | 0.18736 |
| 16 | 389.05 | -2.2464 | 0.087991 | 1.0556 | 0.21998 |

**Table S 10.** The key features were identified using a volcano plot analysis in positive ion mode.

|  | Peaks (mz/rt) | FC | log2(FC) | Raw.pval | -log10(p) |
| --- | --- | --- | --- | --- | --- |
| 1 | 743.5 | 4.6175 | 2.2071 | 4.9054e-08 | 7.3093 |
| 2 | 751.6 | 4.5614 | 2.1895 | 1.9506e-07 | 6.7098 |
| 3 | 839.5 | 4.7283 | 2.2413 | 1.1743e-06 | 5.9302 |
| 4 | 852.45 | 4.7652 | 2.2525 | 2.6082e-06 | 5.5837 |
| 5 | 367.35 | 0.17457 | -2.5181 | 1.0112e-05 | 4.9952 |
| 6 | 811.5 | 4.8491 | 2.2777 | 4.7758e-05 | 4.321 |
| 7 | 397.35 | 0.16605 | -2.5903 | 5.9747e-05 | 4.2237 |
| 8 | 726.5 | 0.14829 | -2.7535 | 0.0024077 | 2.6184 |
| 9 | 899.4 | 0.25215 | -1.9876 | 0.010985 | 1.9592 |
| 10 | 896.5 | 0.24556 | -2.0259 | 0.012977 | 1.8868 |
| 11 | 786.55 | 0.2486 | -2.0081 | 0.01506 | 1.8222 |
| 12 | 487.3 | 8.3493 | 3.0617 | 0.016751 | 1.776 |
| 13 | 348.35 | 0.31755 | -1.6549 | 0.021836 | 1.6608 |
| 14 | 389.3 | 0.33213 | -1.5902 | 0.041732 | 1.3795 |
| 15 | 646.45 | 2.6071 | 1.3825 | 0.049032 | 1.3095 |
| 16 | 788.55 | 0.31997 | -1.644 | 0.055483 | 1.2558 |
| 17 | 805.5 | 0.39977 | -1.3228 | 0.062228 | 1.206 |
| 18 | 895.5 | 0.40723 | -1.2961 | 0.071335 | 1.1467 |
| 19 | 792.55 | 0.29793 | -1.7469 | 0.076521 | 1.1162 |
| 20 | 843.45 | 2.3523 | 1.2341 | 0.083898 | 1.0762 |
| 21 | 376.35 | 0.41829 | -1.2574 | 0.084292 | 1.0742 |
| 22 | 951.35 | 0.42639 | -1.2298 | 0.084704 | 1.0721 |
| 23 | 347.35 | 2.3085 | 1.207 | 0.087745 | 1.0568 |
| 24 | 741.5 | 2.2705 | 1.183 | 0.095579 | 1.0196 |
| 25 | 913.4 | 0.42886 | -1.2214 | 0.096443 | 1.0157 |
| 26 | 619.45 | 0.42838 | -1.2231 | 0.099414 | 1.0026 |

**Table S 11.** Significant features were identified through volcano plot analysis in negative ion mode.

|  | Peaks (mz/rt) | FC | log2(FC) | Raw.pval | -log10(p) |
| --- | --- | --- | --- | --- | --- |
| 1 | 725.5 | 9.7594 | 3.2868 | 1.124e-05 | 4.9492 |
| 2 | 904.55 | 0.27378 | -1.8689 | 2.6398e-05 | 4.5784 |
| 3 | 528.35 | 8.3923 | 3.0691 | 0.00011372 | 3.9442 |
| 4 | 1013.35 | 7.8438 | 2.9716 | 0.00087149 | 3.0597 |
| 5 | 520.2 | 0.28376 | -1.8172 | 0.0011081 | 2.9554 |
| 6 | 496.2 | 0.27156 | -1.8806 | 0.0024573 | 2.6095 |
| 7 | 485.3 | 0.25858 | -1.9513 | 0.0027716 | 2.5573 |
| 8 | 608.4 | 0.27719 | -1.851 | 0.0029671 | 2.5277 |
| 9 | 1127.5 | 3.3403 | 1.74 | 0.029739 | 1.5267 |
| 10 | 1121.4 | 3.3973 | 1.7644 | 0.034602 | 1.4609 |
| 11 | 1191.7 | 2.9848 | 1.5777 | 0.041587 | 1.381 |
| 12 | 1109.6 | 3.0407 | 1.6044 | 0.044677 | 1.3499 |
| 13 | 735.4 | 4.9769 | 2.3152 | 0.053022 | 1.2755 |
| 14 | 392.1 | 0.39388 | -1.3442 | 0.070261 | 1.1533 |
| 15 | 389.05 | 0.44766 | -1.1595 | 0.087991 | 1.0556 |
